# A somatic afterhyperpolarization is driven by ion channel nodes expressed across a polygonal spectrin cytoskeleton

**DOI:** 10.1101/2024.08.08.607230

**Authors:** Giriraj Sahu, Dylan Greening, Xiaoqin Zhan, Wilten Nicola, Ray W. Turner

## Abstract

A spectrin-actin cytoskeleton defines the structure of axons and expression of ion channels that support spike propagation, but it is not known how calcium and potassium channels are organized at the soma to control spike output patterns. The hippocampal pyramidal cell slow afterhyperpolarization (sAHP) is generated by a CaRyK protein complex of Cav1.3 calcium, RyR2, and IK potassium channels reflecting an ER-PM junction. Super resolution imaging and dimension reduction identified a highly organized distribution of CaRyK protein clusters aligned as rows with ∼155 nm periodicity that extended to branchpoints to form a non-rigid lattice-like structure across the soma. All CaRyK proteins proved to align and colocalize with the polygonal spectrin βII cytoskeleton. The data indicate that CaRyK proteins at ER-PM junctions that contribute to the sAHP to control the pattern of spike output are distributed with high precision as functional ion channel nodes across the somatic spectrin cytoskeletal network.

**Graphical Abstract:** 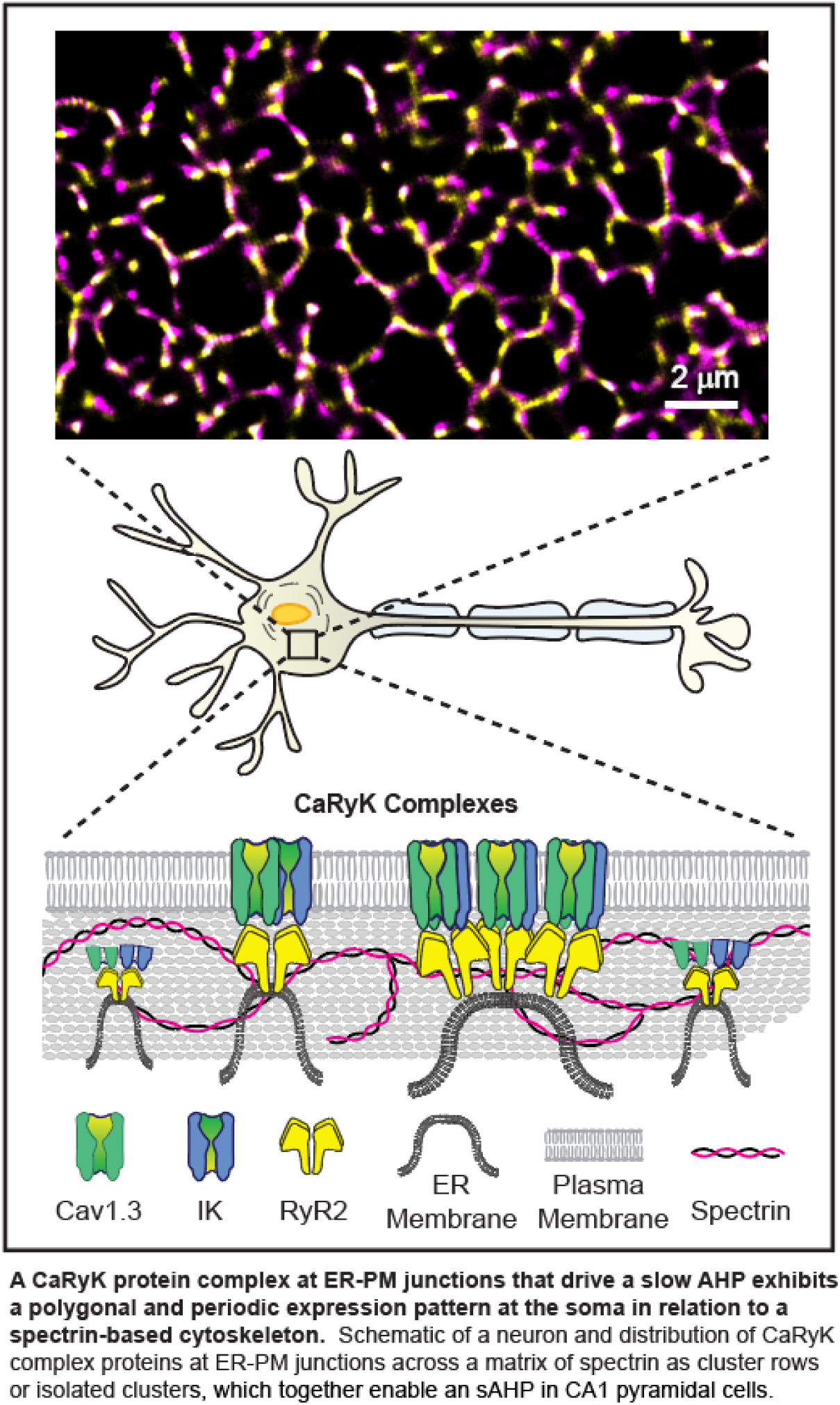

## Introduction

Efficient propagation of a spike from the axon initial segment and down an axon depends on the precise localization of sodium and potassium channels at nodes of Ranvier ^1–4^. By comparison, neuronal somatic membranes express calcium and calcium-gated potassium channels that generate afterhyperpolarizations (AHPs) to pattern spike output into bursts and pauses. In hippocampal neurons a triprotein complex comprised of Cav1.x calcium channels, ryanodine receptor 2 (RyR2), and KCa3.1 (IK) potassium channels (CaRyK complex) form at endoplasmic reticulum-plasma membrane (ER-PM) junctions to generate a slow AHP of several seconds duration ^5–12^. The factors that govern the localization of the CaRyK complex are important as the slow AHP controls spike output in normal and disease states, and circuit function with age ^7,12,13^. Yet the mechanisms by which calcium and potassium channels of the CaRyK complex are spatially organized to control spike output remains to be determined.

It was recently shown that axons exhibit a Membrane Periodic Skeleton (MPS) where lengths of spectrin running parallel to the axon support periodic circumferential bands of actin ^14–20^. Actin forms circular rings at regular intervals along spectrin to create a semi-periodic “1D” pattern ^15,18,19,21,22^. By comparison, a “2D” configuration of spectrin with a polygonal-like organization was reported over regions of the soma or proximal dendritic membranes ^14,16,23^. It is not known if spectrin plays a role in coordinating the somatic expression of a CaRyK complex at an ER-PM junction that contributes to the slow AHP.

We used super resolution STORM-TIRF microscopy to identify protein clusters at the membrane and applied dimension reduction techniques to assess the spatial distribution of the CaRyK complex and its relationship to the cytoskeleton at the soma of hippocampal neurons. Calculations of nearest neighbor distance (NND) and non-negative matrix factorization (NNMF) revealed a coordinated distribution of immunolabel for CaRyK complex proteins across the entire somatic surface. One was a linear series (row) of up to 8 punctate clusters with a periodicity of ∼155 nm that could reach and extend beyond branch points, and a second at distances of ∼650 nm. Together these clusters established a lattice-like structure that was colocalized with that of spectrin βII and actinin linking proteins. Together these data reveal that the CaRyK complex supporting generation of the sAHP exhibits a high degree of spatial organization in relation to a somatic spectrin cytoskeletal lattice.

## Results

STORM-TIRF imaging was used to localize CaRyK complex proteins within 150 nm of the coverslip surface of MAP-2 positive cultured neurons. Images were defined with 10-25 nm localization precision of individual fluorophore centroids. Immunolabels were not assumed to reflect individual ion channels, with a group of fluorophore labels defined by STORM referred to only as clusters ^6,24^. Cluster analysis was performed using masks that included the lateral membrane borders and entire surface of somatic membrane, but not dendritic or axonal processes outside of this region. Nearest neighbour distance (NND) and non-negative matrix factorization (NNMF) calculations were applied to typically >500 clusters per label/cell (675 ± 26; n = 279) and up to 43,000 clusters in up to 55 cells per label for statistical analyses.

### CaRyK complex proteins exhibit a high degree of organization across somatic membrane

STORM-TIRF imaging of immunolabels for proteins that comprise the CaRyK complex revealed discrete labeling of clusters across the entire somatic membrane (***Fig. 1 and 3***). We first focused on cells dual labeled for Cav1.3 and IK immunofluorescence. An extensive distribution of clusters could be detected across large fields of view (***Fig. 1A***) or in magnified ROIs (***Fig. 1B***) as punctate labels in isolation or with discrete overlap of the alternate fluorophore. A distinct pattern was apparent for both IK and Cav1.3 as closely spaced (rows) of clusters (***Fig. 1B***), where ∼50-60% of clusters were contained within rows of 3, ∼20-25% of clusters in rows of 4, and only a few percent in rows of up to 8 clusters (***Fig. 2A***). Rows of clusters could further meet at junction points (***Fig. 1B***) from which 1-2 branches of similarly spaced rows of clusters extended at angles of 70-150 degrees (Cav1.3, 118 ± 2.33 degrees, n = 103; IK, 120 ± 1.59 degrees, n = 101) (***Fig. 2B***). Either IK or Cav1.3 could present as a dominant label, as rows of clusters could be detected for either IK or Cav1.3 (***Fig. 1B***, ROI-3 and 4), with overlap or close association of the other cluster label along a row or at discrete sites that could include the junction point (***Fig. 1B***, ROI-4 and 6). Clusters of immunolabel could also be detected as being physically isolated from cluster rows with or without close apposition of the alternate label (***Fig. 1B***, ROI-5-7).

**Figure 1.**
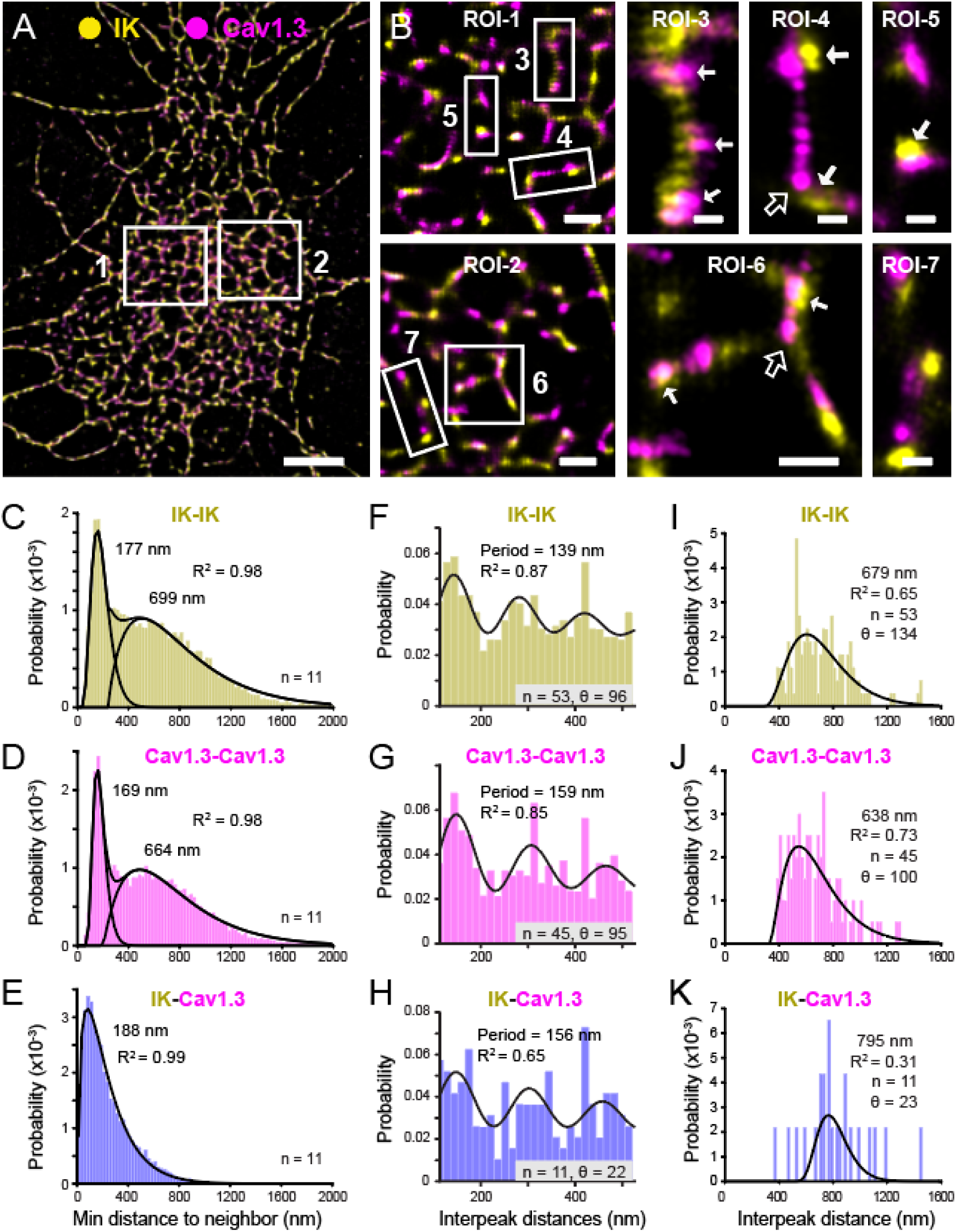
IK and Cav1.3 immunolabels exhibit a coordinated pattern of expression in hippocampal neuronal somata. (**A-B**) STORM-TIRF images of immunolabeled clusters for the indicated proteins detected within 150 nm of the membrane surface. A larger field of view (**A**) and magnified ROIs (**B**) reveal a punctate and organized pattern, with IK or Cav1.3 clusters distinguished as rows of clusters on which the alternate label is closely associated or overlaps at discrete intervals (**B,** ROI-1-4, *solid arrows*). Rows of clusters can reach branch points from which one or two rows of clusters extend (**B,** ROI-4, 6, *open arrows*). In other-cases IK or Cav1.3 clusters are present in isolation or in close association with the alternate immunolabel **(B,** ROI-5, 7). **(C-E)** NND analysis between clusters in dual labeled cells reveals two populations between similar labels (**C-D**) and a right skewed single peak for IK-Cav1.3 (**E**). (**F-K**) NNMF analysis of dual immunolabels that exhibit fluorescence overlap defines a periodic pattern for cluster rows identified by a damped oscillatory model (**F-H**) or isolated clusters (**I-K**). Unless otherwise indicated, in this and all other figures values for fits (**C-E, I-K**) represent the median of gamma distributions. R^2^ values are for the sum of gamma fit distributions (**C-E**), and model periods with R^2^ values (**F-H**). Sample values (n) = cells, θ = features. Scale bars: (**A**) 5 µm; (**B**, ROI-1, -2), 1 µm; (**B,** ROI-6), 500 nm; (**B**, ROI**-**3, 4, 5, and 7), 250 nm. Bin width: (**C-E**) 25 nm, (**F-H**) 15 nm, (**I-K**) 20 nm. See also ***Fig. S1*; *Tables S1 and S2***.

**Figure 2.**
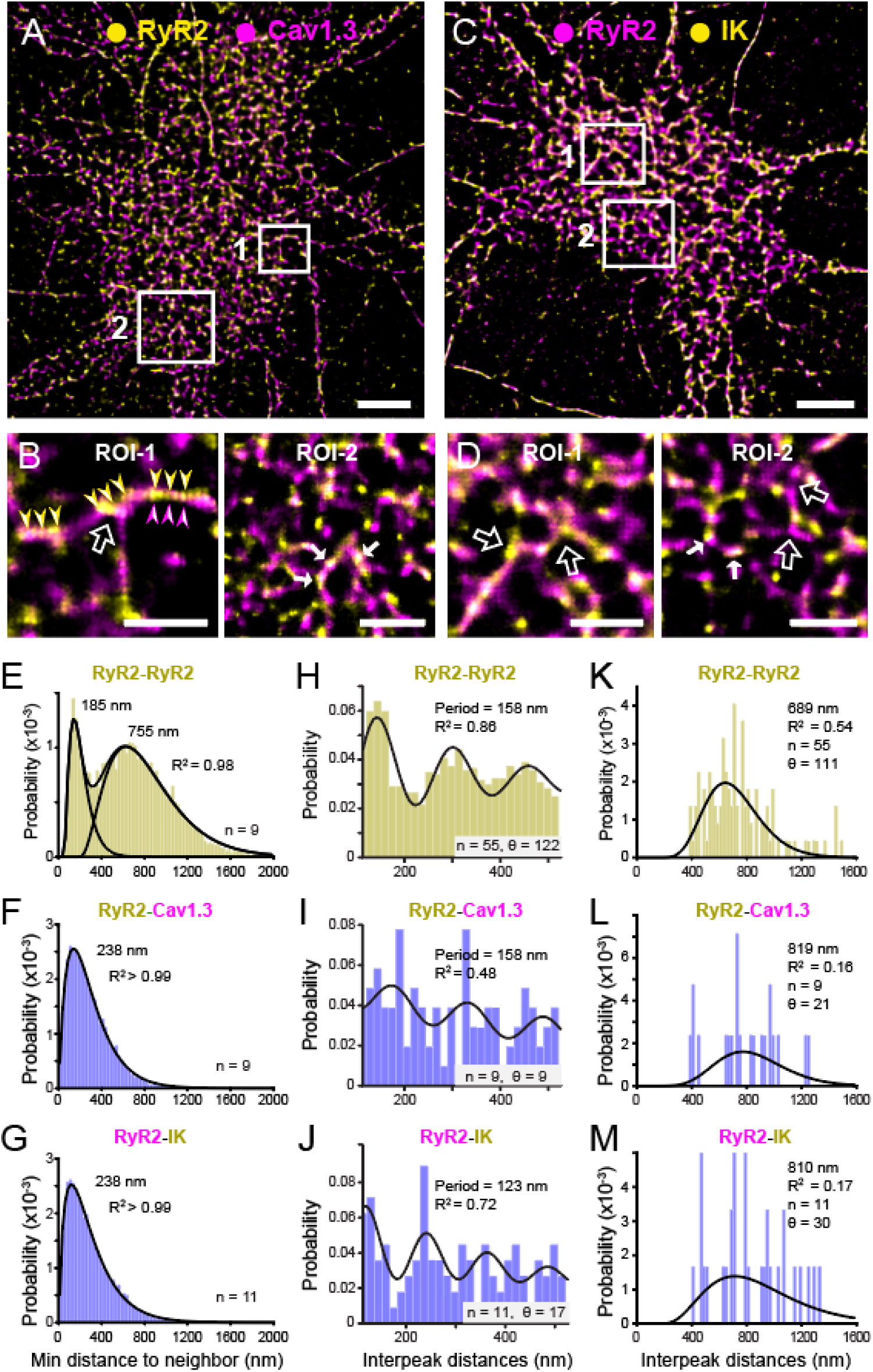
RyR2 exhibits a regular pattern of expression in relation to Cav1.3 and IK. (**A-D**) STORM-TIRF images of immunolabeled clusters for the indicated proteins. Larger field views (**A-C**) and magnified ROIs (**B-D**) show that RyR2 clusters exhibit an organized pattern in relation to Cav1.3 (**A-B**) or IK (**C-D**). RyR2 clusters are distributed in rows (**B,** ROI-1, *arrowheads*) that can reach branch points (**B-D,** *open arrows*) or be positioned in isolation. Cav1.3 and IK labels share these patterns and colocalize with RyR2 as rows of clusters (**B,** ROI-1, *arrowheads;* **B** and **D,** ROI-2, *filled arrows*) or as isolated labels. (**E-G**) NND analysis of dual labeled cells reveals two populations of distance between RyR2 clusters (**E**) and right-skewed single peaks for RyR2-Cav1.3 (**F**) or RyR2-IK (**G**). (**H-M**) NNMF analysis defines a periodic pattern for colabeled cluster rows (**H-J**) or a single distribution for isolated clusters (**K-M**). Sample values (n) = cells, θ = features. Calibration bars: (**A-C**) 5 µm; (**B-D**) 2 µm. Bin widths: (**E-G)** 25 nm, (**H-J**) 15 nm, (**K-M**) 20 nm. See also ***Fig. S1, Tables S1 and S2*.**

**Figure 3.**
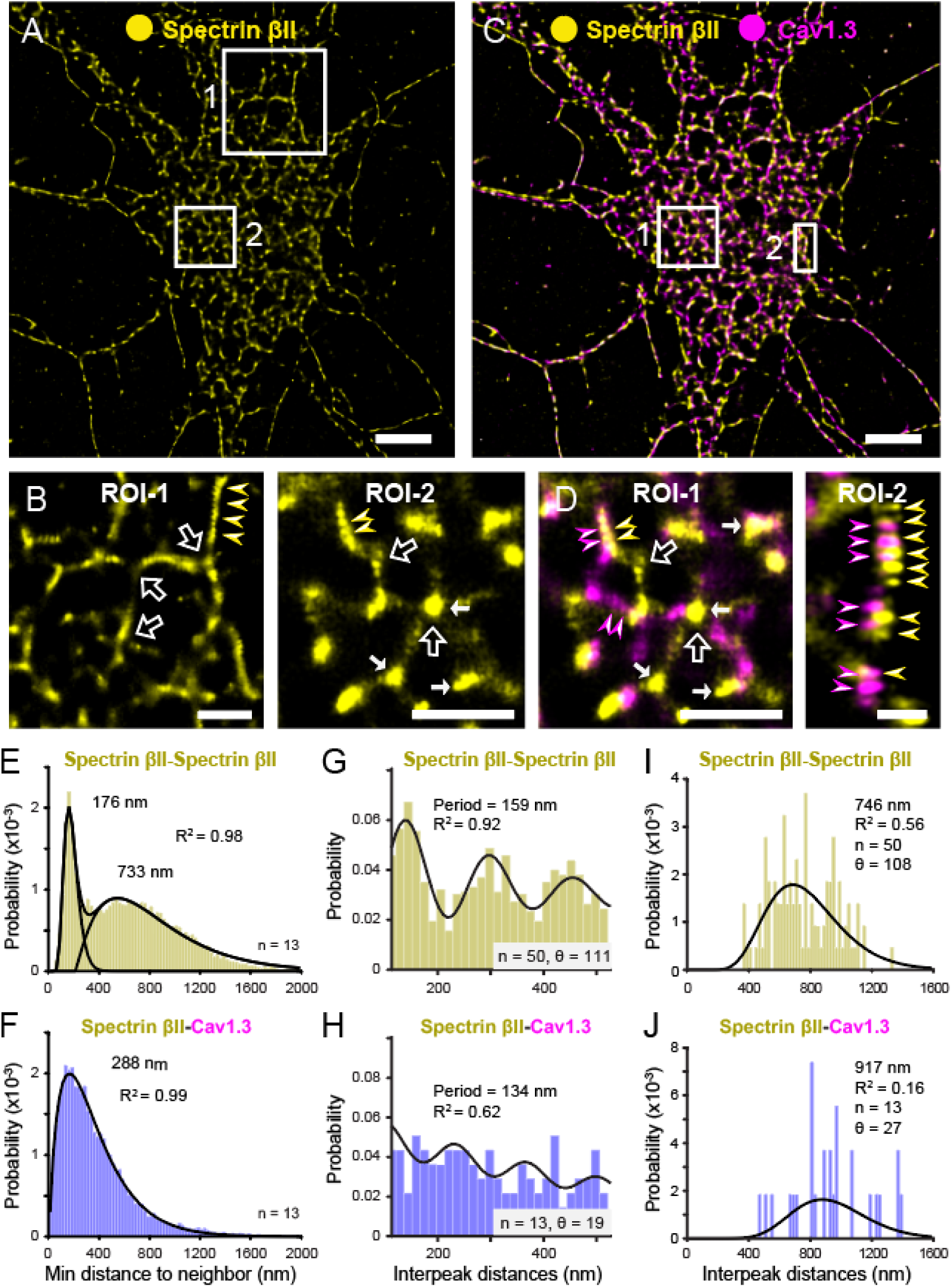
Spectrin βII exhibits a structured pattern associated with Cav1.3. (**A-D**) STORM-TIRF images of immunolabeled clusters for the indicated proteins. Shown are larger fields of view for a cell labeled for spectrin βII (**A** and **B**) and the same cell dual labeled for spectrin βII and Cav1.3 (**C** and **D**). ROIs are magnified in (**B** and **D**), with the same frame of view in (**B,** ROI-2) and (**D,** ROI-1). (**B** and **D**) Spectrin βII labeled clusters organize as rows of clusters (*arrowheads*) that extend to branch points (*open arrows*) or as isolated clusters (*filled arrows*). Dual labeling reveals Cav1.3 clusters that overlap with spectrin βII cluster rows (**C** and **D,** ROI-1, 2, *arrowheads*). (**E-F**) NND in dual labeled cells reveals two populations between spectrin βII clusters (**E**) and one for spectrin βII-Cav1.3 (**F**). (**G-J**) NNMF analysis defines a periodic pattern for cluster rows (**G** and **H**) or a single distribution for isolated clusters (**I** and **J**). Sample values (n) = cells, θ= features. R^2^ value in (**E**) corresponds to the superimposed combined fit for the two populations. Scale bars: (**A** and **C**) 5 µm; (**B,** ROI-1, -2; **D,** ROI-1), 2 µm; (**D,** ROI-2), 500 nm. Bin widths: (**E-F)** 25 nm, (**G-H**) 15 nm, (**I-J**) 20 nm. See also ***Tables S1 and S2*.**

NND calculations identified two inter-cluster relationships for both IK and Cav1.3 (***Fig. 1*, *C and D***). For IK clusters one was fit over a short distance of <500 nm (median of 177 nm), and a second over a larger range of ∼300-2000 nm (median, 699 nm) (***Fig. 1C***). A similar dual distribution of minimal distance was apparent between Cav1.3 clusters up to ∼400 nm (median, 169 nm) and ∼200-2000 nm (median, 664 nm) (***Fig. 1D*, *Table S1***). A plot of the minimal distance between neighboring IK and Cav1.3 clusters instead revealed a single right-skewed histogram of less than ∼1000 nm (median, 188 nm) (***Fig. 1E***). These data are important in confirming a nanometer association between Cav1.3 and IK clusters consistent with colocalization ^6,24^, and two common patterns in the minimal distance between IK and Cav1.3 immunolabels.

### Detecting patterns of protein distribution

The ability to consistently define two patterns of cluster distances between Cav1.3 and IK immunolabels suggests a non-random distribution of CaRyK complex proteins across the somatic membrane. To test this we randomized the centroid positions for each of the CaRyK protein data sets. These tests returned unimodal histograms consistent with spatially random data that were distinct from those of experimental data (***Fig. S1***).

To more fully assess the spatial organization of these labels we conducted NNMF analysis to extract the top 6 detectable features of immunolabel clusters in each cell for analysis (see Materials and Methods, ***Fig. S2-S5***). A key advantage to NNMF is the ability to define features, in an unsupervised fashion, beyond the minimal nearest neighbor for every cluster centroid, without the need to identify the specific orientation of Regions of Interest (ROI) ^25,26^. As a crosscheck of this approach we compared NNMF to the alternate method of autocorrelation applied to predefined ROIs for spectrin βII in the MPS in axons and dendrites and obtained very similar spatial separations of ∼155 nm (***Fig. S4***). It is of interest that these values are somewhat shorter than previous reports of an average value of ∼180-190 nm ^14,15,21^. We consider this to reflect the elasticity inherent to a resting spectrin framework ^27,28^, or potentially the relative length of spectrin subunits incorporated into the developing cytoskeleton at ∼DIV 10-15 ^14,16,29,30^ (see Discussion).

We therefore used NNMF as an objective, automated process to define meaningful patterns of immunolabel clusters at the soma. To identify relationships between colocalized clusters ^6^, we restricted NNMF analysis to only the centroids of clusters identified to exhibit overlap in fluorescence between the two labels (***Fig. S6***).

### CaRyK complex proteins exhibit a regular pattern of expression IK and Cav1.3

NNMF was applied to IK and Cav1.3 clusters to assess the periodicity of rows of clusters as well as isolated clusters. Features extracted by NNMF analysis that reflected a row of clusters were fit by a damped oscillatory model to identify a period for IK clusters of 139 nm (***Fig. 1F***, ***Table S2***). Features with a repeating series of Cav1.3 clusters exhibited a period of 159 nm (***Fig. 1G***). When comparing the relationship between colabeled IK-Cav1.3 clusters, NNMF reported a regularity with a period of 156 nm (***Fig. 1H***). To define potential patterns in the distant (isolated) group identified through NND calculations (***Fig. 1*, *C and D***), we extracted features with peaks separated by distances of 375-1506 nm. This second group of features could be described with a single gamma fit, returning median values of 679 nm for isolated IK-IK clusters, 638 nm for Cav1.3-Cav1.3 clusters, and 795 nm for colabeled IK-Cav1.3 isolated clusters (***Fig. 1I-K***).

These tests established a high degree of shared spatial regularity of IK and Cav1.3 labels as rows of tightly spaced clusters ∼140-160 nm apart or as isolated clusters separated by ∼600-800 nm (***Table S2***). A colocalization of immunolabels for IK and Cav1.3 could be detected for both rows and isolated clusters of distribution.

### RyR2 relationship to Cav1.3 and IK

Previous work established that RyR2s are colocalized with nanometer proximity to Cav1.3 and IK channels as part of a CaRyK complex at ER-PM junctions ^6,7,31,32^. Immunolabeling confirmed extensive colocalization of these labels and a similar spatial organization of RyR2 across the soma (***Fig. 2A-D***). RyR2 clusters could thus be visually distinguished as rows of 3-8 clusters that converged at branch points from which other rows of clusters labeled for RyR2 and/or the counter-labeled protein could extend (mean angle, 116 ± 1.51 degrees, n = 100) (***Fig. 2A-D***). RyR2 clusters could also be identified in isolation with or without overlap of an ion channel fluorophore emission suggestive of colocalization (***Fig. 2*, *B and D***). The latter results are important in emphasizing that RyR2 labeling was not obligatory with Cav1.3 or IK labeling but could exhibit a high degree of co-organization in expression within the 150 nm distance from the surface of the coverslip defined by TIRF imaging.

NND analysis confirmed a separation of RyR2 cluster distances into one group with a nearest neighbor median distance of 185 nm and a more dispersed group (median, 755 nm) (***Fig. 2E***). NND calculations between RyR2 clusters and those for Cav1.3 or IK channels returned right-skewed interval histograms for both RyR2-Cav1.3 and RyR2-IK (medians, 238 nm) (***Fig. 2*, *F and G***). NNMF analysis further revealed a high degree of spatial organization for RyR2, where repeating clusters were fit with a period of 158 nm for both RyR2-RyR2 and RyR2-Cav1.3 clusters, and 123 nm for RyR2-IK clusters (***Fig. 2H-J***). The spatial relationship between isolated clusters for RyR2 was again fit with larger median values for RyR2-RyR2 (689 nm), RyR2-Cav1.3 (819 nm), and RyR2-IK (810 nm) (***Fig. 2K-M***). Altogether these analyses confirmed a close spatial relationship between RyR2 and CaRyK complex ion channels that co-distributed as either rows of clusters or isolated clusters.

### The spatial distribution of CaRyK complex proteins reflects coassociation with the spectrin cytoskeleton

The organized expression of CaRyK complex proteins across the somatic membrane imply the presence of an underlying mechanism for organization. It is known that the spatial distribution of sodium and potassium channels in axons and portions of dendrites is determined by a “1D” spectrin-actin-based MPS ^14–17,19,23,33^. Evidence further points to a “2D” structure formed by spectrin in the somatic region ^14,23^ resembling that of erythrocyte membranes ^34,35^. Several isoforms of spectrin are expressed in neurons ^18,20,29^ with the extent of a 1D structure in dendrites related to the level of spectrin βII ^14,16^. We thus used labeling for the C-terminus of spectrin βII for STORM-TIRF measures of any spatial relationship with respect to CaRyK complex proteins at the soma.

### Spectrin βII and CaRyK complex proteins

Spectrin βII immunolabel could be visualized over the entire region of hippocampal neuron somata (***Figs. 3 and 4***). As found for CaRyK complex proteins, punctate clusters of spectrin βII were detected aligned in rows that reached branch points where 1 or 2 rows of clusters extended at angles with a mean of 118 ±

**Figure 4.**
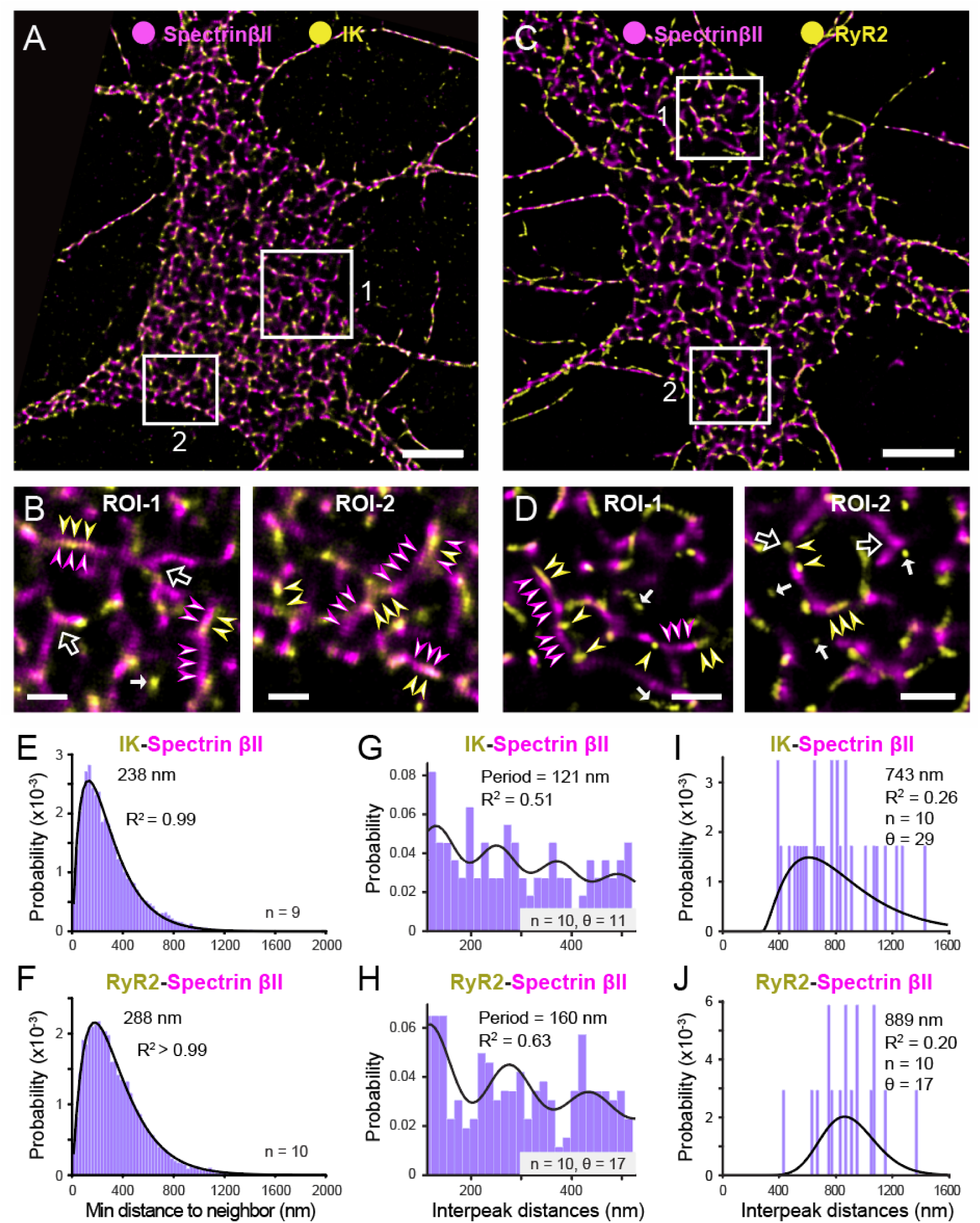
Spectrin exhibits an organized pattern that is closely associated with IK and RyR2 distributions. (**A-D**) STORM-TIRF images of immunolabeled clusters for the indicated proteins. Larger field views (**A** and **C**) with ROIs magnified (**B** and **D**) identify rows of spectrin βII clusters that can reach branch points (*open arrows*) that colocalize or alternate with rows of IK or RyR2 (*arrowheads*). Isolated clusters can be distinguished for IK or RyR2 (**B,** ROI-2, **D**, ROI-1, 2; *filled arrows*). (**E-F**) NND analysis between IK-spectrin βII (**E**) and RyR2-spectrin βII (**F**). (**G-J**), NNMF analysis defines a periodic pattern for cluster rows (**G-H**) or a single distribution for isolated clusters (**I-J**). Sample values (n) = cells, θ = features. Scale bars: (**A** and **C**) 5 µm; **(B** and **D**, ROI-1, -2), 1 µm. Bin widths: (**E-F)** 25 nm, (**G-H**) 15 nm, (**I-J**) 20 nm. See also ***Fig. S7, Tables S1 and S2***.

1.63 degrees (n = 104) or as isolated clusters (***Fig. 3A-D***). In dual labeled cells spectrin βII and Cav1.3 exhibited close alignment that resulted in overlap or close apposition of fluorophore emissions (***Fig. 3*, *C and D***). In some cases the labels were complementary such that labeling of rows of either spectrin βII or Cav1.3 clusters could effectively bridge a row of labels towards an apparent branch point (***Fig. 3D***).

Quantitative analyses revealed many similarities in these patterns of distribution for spectrin βII and Cav1.3 labeling. NND calculations between spectrin βII clusters revealed two populations with short inter-cluster distances of less than ∼400 nm (median, 176 nm), and a second broader distribution up to ∼2000 nm (median, 733 nm) (***Fig. 3E***). The NND histogram between spectrin βII and Cav1.3 clusters was right-skewed with a median of 288 nm, indicating a preferred close spatial relationship (***Fig. 3F***). NNMF analysis of spectrin βII -spectrin βII clusters identified a periodicity of 159 nm in the soma (***Fig. 3G***), similar to that determined for spectrin βII clusters in neural processes (see ***Fig. S4***). Cells co-labeled for spectrin βII and Cav1.3 also identified a regularity in overlapping labels at the soma with a period of 134 nm (***Fig. 3H***). NNMF applied to isolated clusters identified distances of 746 nm between spectrin βII clusters and 917 nm for spectrin βII-Cav1.3 clusters (***Fig. 3I and J***).

Immunolabeling for spectrin βII and either IK or RyR2 revealed similar relationships in organization as that for Cav1.3 (***Fig. 4A-D***). In dual-labeled cells, NND calculations revealed two populations for each of spectrin βII and RyR2 clusters, with an early peak and another with a broader distribution (***Fig. S7***).

By comparison, NND analysis returned right-skewed histograms for spectrin βII-IK (median, 238 nm) and spectrin βII-RyR2 (median, 288 nm) (***Fig. 4*, *E and F***). NNMF analysis of colabeled clusters identified a regular distribution of rows of clusters for spectrin βII-IK (period, 121 nm) and for spectrin βII-RyR2 clusters (period, 160 nm) (***Fig. 4*, *G and H***), and for isolated clusters longer median values for spectrin βII-IK (median, 743 nm) and spectrin βII-RyR2 (median, 889 nm) (***Fig. 4*, *I and J***).

These data are important in revealing that the spatial organization found for each of the three subunits of the CaRyK complex essentially match that of spectrin βII as an indicator of the underlying cytoskeleton.

### Actinin I and II linking proteins

Ion channels in the membrane connect to the actin-spectrin cytoskeleton through intermediate linking proteins ^36^. Key to this are actinins that form extended rod-shaped molecules that cross-link actin filaments, which in turn connect to spectrin ^36^. Previous work has uncovered roles for actinin I and II in controlling the organization of ion channels, including those relevant to subunits in the CaRyK complex ^37–43^.

*Actinin proteins and the CaRyK complex:* Initial tests confirmed a shared spatial organization for spectrin βII and either actinin I or actinin II proteins dual labeled in hippocampal neurons, as confirmed by NND and NNMF calculations (***Fig. S8, S9***). We then examined the spatial relationship between CaRyK complex proteins and actinin I and actinin II. Dual labeled images indicated a very tight association between clusters for actinins and each of Cav1.3, IK, or RyR2 (***Fig. 5A-F***). Immunolabeled clusters were distributed in tightly spaced rows that exhibited strong overlap or alternation in segments of cluster labels between actinin I and Cav1.3, actinin II and IK, and actinin II and RyR2 (***Fig. 5B, D, and* *F***). Indeed, the extent of colabeling of actinin II cluster rows with IK or RyR2 provided some of the clearest indications of expression consistent with an underlying spectrin-based architecture ^37,41^. NND analysis restricted to clusters of actinin and CaRyK complex proteins presented interval histograms that were strongly right skewed, with median values between different pairs of clusters ranging from 138-213 nm (***Fig. 5G-I***).

**Figure 5.**
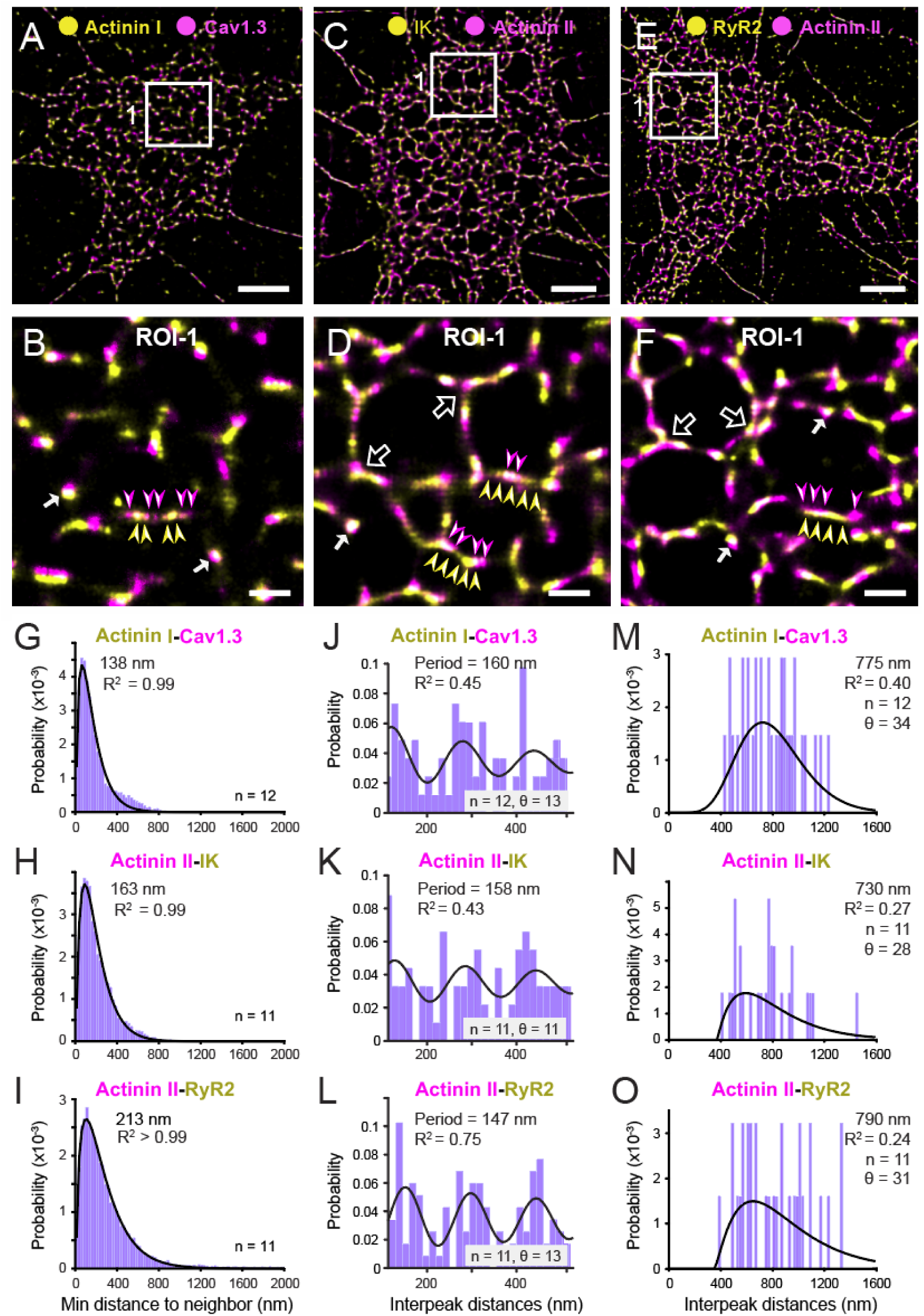
Actinin I and actinin II distributions are aligned with CaRyK complex proteins. (**A-F**) STORM-TIRF images of immunolabeled clusters for the indicated proteins. Shown are dual labeled im-ages for the indicated CaRyK complex proteins and actinin isoforms. Larger field views (**A, C, E**) and associated ROIs (**B, D, F**) identify clusters that align in rows that either overlap or alternate (*arrowheads*) with actinin isoforms. Cluster rows can extend to branch points (*open arrows*), colocalize (*solid arrows*), or alternate to extend or bridge rows of labels (*arrowheads*). (**G-I**) NND analysis between actinin I - Cav1.3 (**G**), actinin II - IK (**H**) and actinin II - RyR2 (**I**). **(J-O**) NNMF analysis defines a periodic pattern for cluster rows (**J-L**) or a single distribution for isolated clusters (**M-O**). Sample values (n) = cells, θ= features. Scale bars: (**A, C, E**) 5 µm; (**B, D, F**), 1 µm. Bin widths: (**G-I)** 25 nm, (**J-L**) 15 nm, (**M-O**) 20 nm. See also ***Figs. S8 and S9, Tables S1 and S2*.**

NNMF analysis also confirmed a periodicity of 147-160 nm between colabeled actinin - CaRyK complex proteins, and for isolated clusters of 730-790 nm (***Fig. 5J-O***).

### Correspondence between the spatial organization of CaRyK complex and cytoskeletal proteins

One of the most striking parallels in the spatial organization of cytoskeletal and CaRyK complex proteins was in colabeling rows of clusters that reached or extended beyond branchpoints. To further compare these parameters, we examined rows of clusters at high resolution in up to 5 randomly selected ROIs of 600 X 600 pixels / cell drawn from 13-55 cells per label (***Fig. 6A, B***). Up to 388 rows for CaRyK complex proteins and spectrin βII were defined across a selection of up to 275 ROIs and 55 cells (***Fig. 6A***).

**Figure 6.**
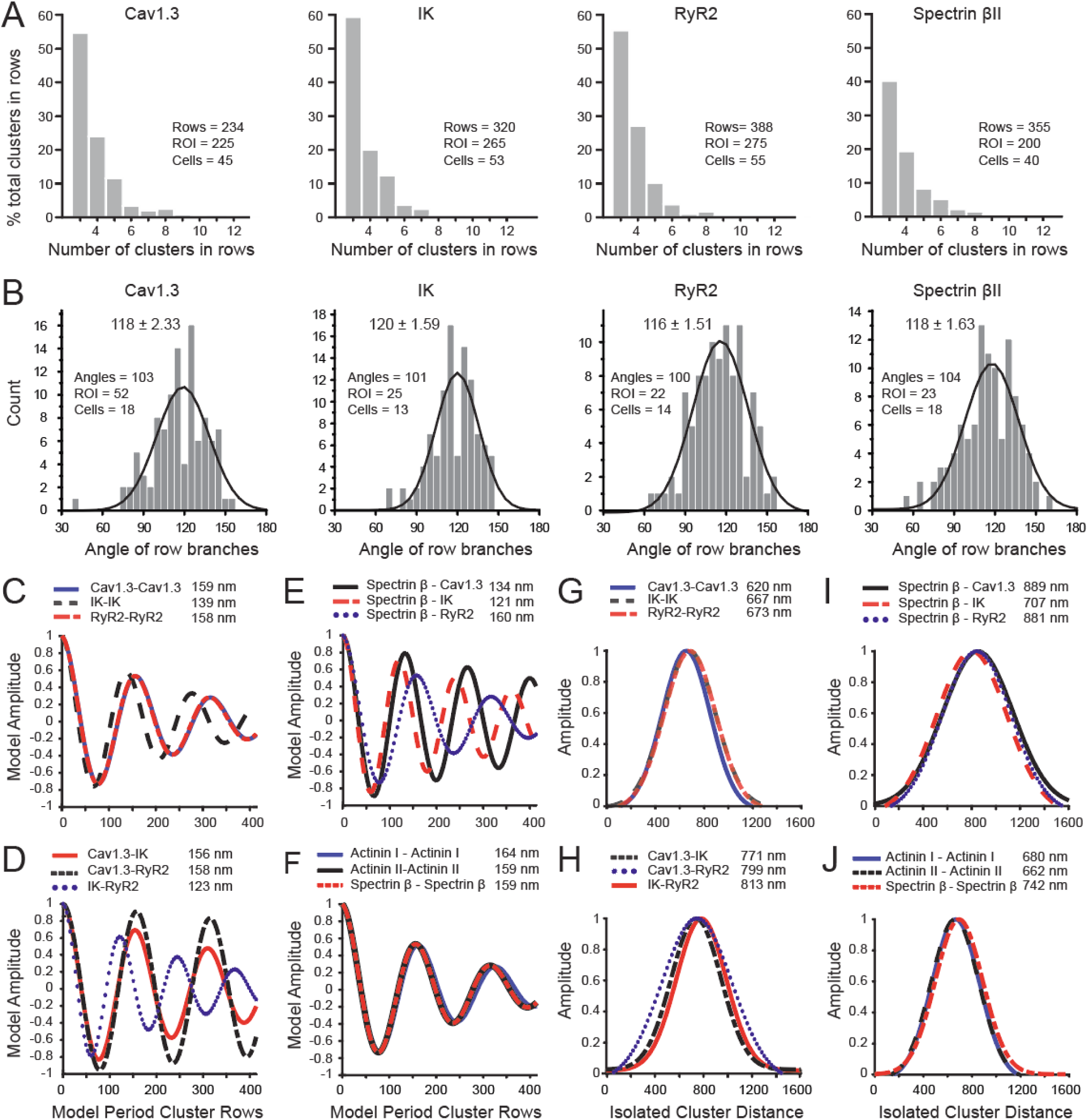
Correspondence between the spatial organization of CaRyK complex proteins and spec-trin. (**A**) Plots of the number of clusters aligned in rows of at least three clusters. (**B**) Plots of the angles at branch points of cluster rows. Bin widths: 5 degrees. (**C-J**) Comparison of model fits for cluster rows (**C-F**) and isolated clusters (**G-J**). The amplitude and peak time of damped oscillatory model fits are normalized to the clusters used in NNMF (first crest of fit, 0 nm) (**C-F**) and gaussian centers (**G-J**) are normalized only to amplitude (**G-J**). Fits are from Fig. 1F**-K**, Fig. 2H**-M**, Fig. 3G**-J**, Fig. 4G**-J**.

Comparing the number of clusters per row returned very similar histogram distributions for each of the CaRyK complex proteins. Approximately 50-60% of all clusters were contained within rows of 3 clusters, with a decrease in number to ∼20-25% of clusters in rows of 4, and to only a few percent in rows containing up to 8 clusters (***Fig. 6A***). By comparison, ∼40% of spectrin βII clusters were contained in rows of 3 clusters, with a similar decrease in number from 4-8 clusters per row (***Fig. 6A***).

We further compared the angles formed at branch points of rows for CaRyK complex proteins and spectrin βII. Measurements were drawn for rows of least 3 consecutive clusters that reached a point where a second row of clusters extended with a shift in orientation. Approximately 100 branch points were measured for each label across 22-52 ROIs randomly selected from 13-18 cells (***Fig. 6B***). In each case, branch points were formed at angles that ranged between 60-150 degrees centered on a mean value of ∼118 degrees (***Fig. 6B***). No statistical difference was found between any of the distributions of angles for rows of CaRyK proteins or spectrin βII clusters (***Fig. 6B; Table S3***).

Additional comparisons were made for the fits obtained by the damped oscillatory model for cluster rows and gaussian fits for isolated clusters (***Fig. 6C-J***). The close spatial relationship between CaRyK complex proteins was emphasized for the model fits for rows of Cav1.3-Cav1.3 and RyR2-RyR2 that entirely superimposed (***Fig. 6C***). The fit for the period of IK-IK cluster rows overlapped these distributions, although the period was ∼19 nm shorter (***Fig. 6C***). When comparing the spatial pattern for CaRyK proteins that exhibited overlap in emissions, the model fits for Cav1.3-IK and Cav1.3-RyR2 showed closely aligned peaks with periods of ∼157 nm (***Fig. 6D***). By comparison, the model fit for the period of IK-RyR2 clusters presented with a shorter relative period in which crests peaked ∼34 nm shorter than for Cav1.3-IK and Cav1.3-RyR2 labels (***Fig. 6D***). The spatial regularity revealed by NNMF for CaRyK complex proteins in relation to spectrin also showed a close correspondence (***Fig. 6E***). This was the case for model fits of the period of rows of clusters between spectrin βII and Cav1.3 (134 nm) and spectrin βII and IK (121 nm) (***Fig. 6E***). The period of cluster rows between spectrin βII and RyR2 (160 nm) conformed somewhat less to the fits for Cav1.3 and IK, with a period ∼30-40 nm longer (***Fig. 6E***). By comparison, the model fits for the relationship between rows of clusters labeled for actinin I or actinin II with respect to spectrin βII almost perfectly superimposed (∼160 nm) (***Fig. 6F***), emphasizing the close spatial relationship between these linking proteins and spectrin βII.

It is interesting that the damped oscillatory model indicated a shorter relative period for IK clusters compared to Cav1.3 or RyR2 (***Fig. 6C, D***). This could suggest a greater number of IK clusters or a spatial organization at shorter relative intervals. However, the total number of centroids detected for IK vs Cav1.3 per cell was not statistically different (IK clusters; Cav1.3 clusters; p = 0.1413; 2-sample Kosmolov-Smirnov test). The shorter period for IK clusters might instead reflect an association between IK channels and proteins other than the CaRyK complex ^6,44^, decreasing the relative spacing between IK clusters. RyRs are also known to associate with several other potassium channels (BK, SK, Kv4) ^45–47^ that could be reflected in the relative spacing of IK-RyR2 clusters. Finally, RyR2 can also be expected to be localized in the ER for functions other than associating with the CaRyK complex.

Comparison of the gaussian fits for the spatial distribution of isolated clusters of CaRyK complex proteins were very consistent in revealing a peak inter-cluster distance between each class of protein of ∼650 nm (***Fig. 6G***), and a close alignment of fits for isolated clusters of CaRyK complex proteins with overlapping signals of ∼800 nm (***Fig. 6H***). Similarly, gaussian fits for the distance between isolated clusters for overlapping emissions of spectrin βII and both Cav1.3 and RyR2 superimposed with values of ∼800 nm, with a slightly shorter fit of 707 nm for spectrin βII and IK (***Fig. 6I***). As noted for actinin protein relationships with nearest neighbor distances to CaRyK complex proteins, the gaussian fits for actinin proteins or spectrin βII were of shorter but almost identical peak distances of ∼700 nm (***Fig. 6J***).

## Discussion

The mechanisms that define the expression of membrane ion channels and the protein-protein interactions needed to create complex patterns of neuronal output has been a long-standing area of investigation. It has now been established that spike propagation down an axon is enabled by a spectrin-actin MPS that creates a highly ordered structure to localize sodium and potassium channels at nodes of Ranvier ^15,18,19,21,22^. The pattern and frequency of spikes are instead controlled by ion channels at the *soma* that use changes in calcium concentration to modify their level of activation. Specialized ER-PM junctions where the ER reaches to within 10-30 nm of the somatic plasma membrane can coordinate activity of ion channels and internal calcium release sites to control cell excitability ^48,49^. One representative assembly at ER-PM junctions is the CaRyK complex that contributes to generating a slow AHP in hippocampal neurons that shapes the pattern and frequency of spike output ^5–7,11,50^. The current study reveals that proteins of the CaRyK complex are not randomly distributed at the soma. Rather, the CaRyK complex exhibits a coordinated distribution across the somatic surface consistent with a pattern defined by a spectrin-based cytoskeletal matrix ^14^.

### Patterns of organization of the CaRyK complex

STORM-TIRF images revealed that CaRyK proteins exhibit a polygonal organization at the soma that was defined by NND and NNMF analyses as reflecting a periodic distribution. We thus identified rows of immunolabels composed of ∼3-8 clusters spaced ∼155 nm apart for each of the CaRyK complex proteins. The rows of clusters further reached branchpoints from which additional rows of clusters extended at internal angles of 60-150 degrees (mean of 118 degrees) that could effectively close loops in the branching structure. A second prominent pattern for CaRyK proteins and spectrin βII was in isolated clusters with a peak distance of ∼600-800 nm. Furthermore, each of these measures were consistent between CaRyK complex proteins, actinin linking proteins, and spectrin βII (***Tables S1-S3***).

The MPS of axons, the AIS, and some dendrites have been shown to exhibit a “1D” pattern of longitudinally oriented spectrin repeats and circumferential bands of actin at regularly spaced intervals ^14–17,19,20,51,52^. The somatic surface of neurons has instead been reported to exhibit a 2D structure of spectrin labeling that undergoes an increase in complexity over time to reach ∼45% of cell surface area by DIV 28^14^. The current work employed NNMF as a dimension reduction technique ^25,55^ to objectively define a lattice-like architecture assembled by spectrin, presumably to support the larger surface area of neuronal somata. This structure is then reflected in the pattern of actinin I and II that act as linking proteins to CaRyK complex proteins at ER-PM junctions. The distance between spectrin repeats at the soma using an antibody against the C terminus of spectrin βII was fit by an oscillatory model at ∼159 nm (R^2^ = 0.92). Our use of NNMF on a select number of MAP-2 negative (presumed axons) or MAP-2-positive (presumed proximal dendrites) processes identified mean values of 148 nm (R^2^ = 0.90) for axons and 157 nm (R^2^ = 0.84) for dendrites (***Fig. S4***). These values compare to reported distances between spectrin repeats in the MPS of unmyelinated axons of 182 ± 16 nm (range of 150-220 nm) ^15^ and for myelinated axons of 180 ± 35 nm (range of 80-240 nm) ^17^. This range of values would be consistent with the known elasticity of spectrin ^19,27,35,53,54^, and validates NNMF as an objective procedure to define the periodicity of proteins ^25,55^. The current work is thus important in establishing that the spectrin cytoskeleton provides a framework to organize the expression pattern of an ion channel complex positioned as part of ER-PM junctions at the soma, a pattern that may be prevalent in the distribution of other somatic ion channels.

### Functional role of a periodic distribution of the CaRyK complex

Previous work has shown that IK channels that are part of the CaRyK complex contribute substantially to the sAHP of CA1 hippocampal pyramidal cells ^5–7,11^. A wealth of data on sAHP properties ^7^ provides a plausible explanation for why the CaRyK complex organization exhibits such a high degree of periodicity across the somatic membrane. One could anticipate an advantage to localizing the CaRyK complex at specific spatial distances to reduce energy expenditure and generate a consistent AHP. Indeed, aberrant modulation of the sAHP has been shown to compromise circuit function with age or promote temporal lobe epilepsy ^10,13,56^. Interestingly, previous work reveals a strong functional coupling between L-type calcium channels and shows that single stimuli can activate sAHP channels in CA1 pyramidal cells ^57^ but typically do not evoke a prominent sAHP ^7^. Instead, activating sAHP channels with a high Po depends on a minimum threshold of membrane activity, such that short trains of inputs (50 Hz, 5 stimuli) rapidly recruit “sAHP” and IK channels ^5,50^. Once evoked by a stimulus train, the sAHP further exhibits a characteristic delay to onset and peak amplitude ^50,58,59^.

The minimum stimulus threshold for sAHP activation could reflect properties of the calcium sources and/or their ability to interact with the IK calcium sensor. We note that previous reports determined that all members of the CaRyK complex exhibit protein-protein interactions at the nanodomain level (<10 nm distance) ^6^, ensuring proximity of Cav1.3 and RyR2 to calmodulin as the calcium sensor for IK channels^60^. The level of increase in calcium concentration obtained upon activating calcium channels will depend on the number of channels ^61^, single channel conductance ^61^, kinetics ^8,61^, facilitation of calcium channel activity ^8,59,62^, and secondary activation of RyR2-mediated calcium release ^6,63,64^. Together these define radial increases (“dome”) in internal calcium defined either as a calcium nanodomain within 20 nm of the channel, or a calcium microdomain of lower calcium concentration with radius of 100-200 nm ^65^. To our knowledge, modeling of the calcium increase mediated by RyRs has not been reported for neurons, although a similar nanodomain and microdomain configuration is assumed.

In pyramidal cells, several factors contribute to an increase in calcium influx to shape its spatial and temporal influence. First, both Cav1 channels and ryanodine receptors can be expressed in clusters^66,67^, and Cav1 channels can exhibit a stimulus-dependent aggregation ^68^. Repetitive stimulation in CA1 pyramidal cells also evokes a frequency-dependent and delayed facilitation of Cav1.3 channels that is imparted on their activation of IK channels ^8,59,62^. Cav1.3 calcium influx further recruits ryanodine-evoked calcium increases of longer duration ^67,69^. The relative delay to onset and peak of the sAHP following repetitive stimulation is thus thought to reflect at least the properties of Cav1.3 channels and ryanodine receptors.

Given these considerations, the periodic localization of CaRyK complexes at ∼155 nm could be ideal for functional activation of the sAHP. Depolarization will open Cav1.3 channels to form a calcium nanodomain of ∼20 nm which would account for the functional coupling between single L-type and sAHP channels ^57^, but falls well below the 155 nm distance to the next CaRyK complex and ER-PM junction. During repetitive stimulation calcium influx will increase due to the factors discussed above, promoting the development of a calcium microdomain of ∼200 nm radius that will overlap with neighboring ER-PM junctions. The stimulus-dependent threshold for activating an sAHP at the whole cell level may then reflect a sequential and temporal recruitment of a minimal number of IK channels at neighboring ER-PM complexes as the calcium increase spreads spatially to create a calcium microdomain. Such a process could further account for why sAHP amplitude is directly related to the number and frequency of stimuli, and evoked following a repetitive stimulus with a delayed onset and peak for activation. Expression of the CaRyK complex in relation to the spectrin cytoskeleton may thus coordinate activation of the sAHP across somatic membrane.

### ER-PM junctions as ion channel nodes

ER-PM junctions have previously been shown to coordinate functional outcomes that include endo- and exocytosis, calcium rebalancing, membrane protein delivery, lipid transfer, and excitation-transcription coupling ^32,70–74^. ER-PM junctions can also coordinate the expression of calcium channels at the plasma membrane and RyRs in the ER ^6,31,48,49,75^. The combination of Cav1.3-RyR2-IK proteins studied here as component elements of ER-PM junctions are necessary for the generation of a slow AHP ^6–8^. Spikes near the soma can thus regulate firing rate by activating calcium increases to produce a slow AHP of up to secs duration. The current work highlights how the CaRyK complex exhibits a lattice-like organization that follows the spectrin-based architecture. The data would then support a role for ER-PM junctions ^48^ as ion channel nodes that colocalize calcium- and calcium-gated potassium channels at the soma with a spatial distribution defined by the spectrin cytoskeleton. When assembled and distributed in this manner, calcium entry and/or internal calcium release can trigger the calcium-activated potassium channels required to regulate the frequency and pattern of spike output. The MPS of axons is instead tailored to organize sodium and potassium channels for the efficient propagation of spikes from the soma to postsynaptic structures. It needs to be considered that Kv2.1/2.2 potassium channels have been shown to control assembly of a host of proteins at ER-PM junctions that include calcium channels and RyRs ^31,48,49,67,71,74–76^. At this time we do not know the extent to which Kv2.x channels are responsible for organizing either of the patterns observed here for subunits of the CaRyK complex, but anticipate that future work will show a similar role for the spectrin cytoskeleton in organizing the expression pattern of many other ion channels.

### Limitations of the current study

We note that the extent to which we could define the full structure of a spectrin-based cytoskeleton was limited by several factors. For one, neurons are known to express tetramers of *αα* and different β spectrin subunits ^18–20,77^. Our labeling and NNMF calculations centered on the spectrin βII C terminus and were restricted to dual labeled protein pairs exhibiting overlapping fluorescent emissions, reducing the total number of clusters that might align in a given sequence. For these reasons the full extent of a spectrin-based polygonal structure was likely underestimated.

## Methods

### Experimental Design

The objective of this study was to use super resolution STORM-TIRF imaging of immunolabels for a CaRyK protein complex in relation to markers of the cytoskeleton at the soma of hippocampal pyramidal cells in dissociated cultures. Dimension reduction approaches were used to objectively quantify the physical relationship and patterns of CaRyK protein immunolabels in relation to spectrin.

All materials and supplies with catalog numbers, software, and site of data repository are provided in ***Table S5*.**

### Animals

Timed pregnant Sprague Dawley rats were obtained (Charles River) and maintained for at least 7 days after shipping according to the guidelines of the Canadian Council of Animal Care and experimental protocols approved by the University of Calgary Animal Care Committee.

### Dissociated hippocampal cell cultures

P0-P1 male or female rat pups were used to prepare low-density dissociated cultures of rat hippocampal neurons ^6,8^. P0-P1 rat pups were anesthetized on ice for 3-5 min and hippocampi dissected and dissociated by papain treatment followed by titration with progressively smaller bore glass pipettes. After each titration step the supernatant containing the cells was passed through a 70 µm separation filter (Corning, Cat. 07201431) into a 50 ml tube. Cells were centrifuged at 300 x g for 10 min and the supernatant completely aspirated and resuspended in MACS buffer (cat#130-091-221) (diluted 1:20). Unlabelled hippocampal neurons were collected using a MiniMACS™ Separator (130-042-102). Prior to collection, cells were incubated with anti-rat CD11b/c antibody (Cat#130-105-634) conjugated to magnetic microbeads.

After microglia depletion, cells were incubated with GLAST (ACSA-1)-biotin (Cat#130-095-826) according to manufacturer instructions (Miltenyi Biotec, CA) (***Table S5***) ^78^. Collected unlabeled neurons were diluted in basal neural medium (BME) supplemented with 1 mM Na-pyruvate, 2 mM L-glutamine, 10 mM HEPES, 1% B-27 supplement, 5% FBS, 0.6% glucose, and 1% penicillin-streptomycin for correct plating density. The cell suspension was then plated on laminin and poly-l-lysine-coated glass coverslips in 12 well culture dishes at 3500–5300 cells/cm^2^. Cells were grown on 18 mm No 1.5 coverslips (Electron Microscopy Sciences, PA) at 37 °C for three days in BME, followed by exchange with BME lacking FBS. Culture medium was refreshed with BME lacking FBS every three days. Cultured hippocampal neurons at 10-15 days *in vitro* (DIV 10-15) were used to obtain images of isolated neurons to further reduce the risk of glial membrane envelopment that could interfere with imaging neuronal proteins adjacent to the coverslip surface.

### Immunocytochemistry

At 10-15 days *in vitro* (DIV) coverslips with cells were washed with phosphate-buffered saline (PBS) and fixed in 4% paraformaldehyde (PARA) (Electron Microscopy Sciences, ON) in PBS for 10-15 min. After 3 washes in PBS cells were incubated with a blocking medium consisting of (5% (w/v) bovine serum albumin (Vector Laboratories) and 0.2% (v/v) Triton X-100 (Sigma Aldrich) in PBS for 45 min at RT (∼22°C) for all subsequent treatments (***Table S5***). Cells were incubated with primary antibodies in a blocking medium for 45 min at RT. After five 10 min washes in blocking medium cells were treated with the secondary antibodies for 45 min at RT. After three rounds of 10 min washes in blocking solution cells were washed once in PBS and post-fixed with 4% PARA and 0.1% glutaraldehyde in PBS for 10 min followed by three 10 min washes in PBS. TetraSpek™ beads (1:2000, 100 nm diameter, T7279, Thermofisher) were added to coverslips overnight at 4°C in PBS followed by three 5 min washes in PBS to remove loose beads.

### Direct Stochastic Optical Reconstruction Microscopy (dSTORM)

dSTORM imaging was used together with TIRF illumination to localize CaRyK complex proteins within 150 nm of the coverslip surface of MAP-2 positive cultured neurons. Imaging cocktail was prepared fresh from 1X PBS, 10% (w/v) glucose, 10 mM β-mercaptoethanol in an oxygen-scavenging GLOX solution containing 0.5 mg/ml glucose oxidase and 40 μg/ml catalase (Sigma-Aldrich). 10X GLOX solution was prepared as a stock and stored in aliquots at −20°C. On the day of imaging an aliquot of GLOX solution was used to prepare the dSTORM imaging buffer. Imaging cocktail was placed into a concavity slide, covered with the immunolabeled 18 mm coverslip and sealed with VALAP (equal parts vaseline, lanolin and paraffin). Images were obtained using a Quorum Diskovery Flex microscope equipped for TIRF illumination (150 nm depth) using a HC Plan Apo 63X/NA 1.47 oil immersion objective and Andor iXon Ultra 897 EMCCD camera. Fluorophores were first photo switched into a temporary dark state using strong intensity laser illumination (laser power: 561 nm, 55 mW @ 100%; 647 nm, 53 mW @ 100%). Subsets of active blinking fluorophores were then imaged at a lower laser intensity (70-95% excitation) using a 33 Hz frame rate. The 405 nm laser line was adjusted to obtain a balanced reactivation of Alexa Fluor-647 or Cy-3 dyes during image acquisition of 5000-8000 frames per sample.

Image reconstruction was carried out as previously described ^6^ using a ThunderSTORM plugin (Fiji, ImageJ). A calibration factor of 150.5 nm / pixel was obtained under our imaging conditions after rigorous calibration using a slide mounted Supracon nanoscale standard linewidth/pitch target (Part number Kali2020_2_F5; Supracon, Germany) (Live Cell Imaging Facility, University of Calgary). The acquired images were filtered using the wavelet filter function of ThunderSTORM with default values (beta spline order of 3 and a scale of 2), for determining signal-to-noise threshold for the datasets. The camera parameters used were pixel size: 150.5 nm; photoelectrons per A/D count: 8.98; base level A/D count: 200; EM gain: 300. Afterward, the point spread function of the imaged fluorophore localizations were detected with the standard gaussian filter method and fitted with a weighted least square fitting using the subpixel localization fitting procedure of ThunderSTORM. Only the localizations falling under a standard deviation of the fitting distance, representing sigma >50 and <500 were taken into further analysis. Any mechanical drift over the timeframe required for imaging over 15-20 min (0 to <500 nm) was drift corrected using the cross correlation function of ThunderSTORM, with any cases approaching 700 nm excluded from analysis. Correct image reconstruction and alignment for dual labeled images were assured using the X-Y coordinates of 3-6 100 nm fiducial beads (T7279, Thermofisher) as reference points to align signals with an affine/linear registration function using Localization Microscopy Analyzer (LAMA) ^79^. X and Y positional coordinates and intensity information of point localizations were analyzed using the morphological cluster analysis pipeline of LAMA software to identify the coordinates of protein clusters with a minimum cluster radius of 25 nm.

### Image Analysis

Extensive work on the pattern of actin and spectrin labeling in axons and dendrites has defined a structural entity referred to as an MPS ^14–20^. Obtaining a measure of the periodicity of proteins along the MPS was obtained using autocorrelation on ROIs that defined the linear projection of given axons or dendrites. The issue is more complicated in somatic regions where even closely spaced immunolabel clusters could project in any x-y coordinate. For this reason both NND and NNMF were used to systematically search through pixels in an ROI that included the lateral borders and entire cytoplasmic region of the soma defined by MAP2-GFP labeling.

STORM analysis of spatial relationships was carried out on either single immunolabels or on centroids reflecting an overlap of fluorophore emissions by calculating the geometric mean (***Fig. S6***). While the extraction of centroids with overlapping pixels served to eliminate single labeled clusters, we note that it also had the effect of underestimating the total number of co-associated clusters that appeared to be in immediate apposition, if only through a function of parallax of vertically aligned proteins. For example, with STORM imaging of cells dual labeled for spectrin βII and Cav1.3, the total number of spectrin βII centroids was 6,741. The process of extracting spectrin βII clusters that overlapped with Cav1.3 clusters reduced the detected spectrin βII centroids by 53% to a total of 3,191. Despite these reductions in centroids detected, NNMF analysis typically applied to 8-13 dual labeled cells with 900-5,100 centroids that generated 9-22 features for clusters associated in rows and 14-34 features for isolated clusters. Analysis of the percentage overlap of colabelled CaRyK protein clusters as determined by the portion of pixels with intensity higher than a threshold of 0.2 in both channels is shown in ***Table S4***.

### Nearest Neighbour Distance (NND)

Intensity information and close proximity of individual fluorophores of the ThunderSTORM/LAMA reconstructions were used to create a mask and identify immunolabeled clusters using the standard morphological cluster analysis (MCA) module of LAMA ^6,24,79^. Cluster centroid information obtained from MCA analysis was used to determine nearest neighbour Euclidean distances in MATLAB ^6,24^. Distributions were computed for each immunolabel cluster set (eg. Cav1.3-Cav1.3, Cav1.3-RyR2) and fit with either a single gamma distribution or a gamma mixture model using the MATLAB function *lsqnonlin* to define central tendency of the sub-populations in the distributions ^80^. Mixing portions are the integral of each gamma distribution in comparison to the integral of the gamma-mixture model. To assess the spatial randomness of clusters, the NND distributions were further compared to that of the same number of centroids with randomized positions and histograms assessed with the Kosmolgov-Smirnov test in MATLAB (***Fig. S1***).

### Non-negative matrix factorization (NNMF)

NNMF is a computational algebraic tool for reducing dimensionality in data ^25,26^. As a form of singular value decomposition, NNMF generates two matrices reflecting a collection of ‘features’ and weights’, of which the product approximates the input data. A strength of NNMF is its capacity to define patterns within a dataset when data has been standardized. Ultimately NNMF provides a means to compress hundreds of standardized images and reduce them into a handful of ‘feature’ images (for reviews see ^25,55^).

A brief summary of the process for NNMF analysis used here is presented in ***Fig. S2-S5*** for the case of spectrin βIIC imaged by STORM-TIRF in neural processes external to the soma, with parameters used for extracting features from somatic labels applied to these images. First, STORM-TIRF images confirmed that the resolution of our system was sufficient to identify the repeating domains of spectrin βII in both MAP2 negative (presumed axons) and MAP2 positive (presumed dendrites) of cultured neurons consistent with an MPS structure. Images were prepared for NNMF analysis using a MATLAB script which normalized image intensities and identified centroids through image segmentation using the *regionprops* function. To standardize pattern detection for clusters initially oriented in any x-y direction, ROIs centered around every centroid position were stored as 2D 600-X-600 pixel (4.5 X 4.5 µm) images and rotated so the nearest centroid was oriented to the left (***Fig. S2***). The MATLAB function *NNMF,* from the ‘machine learning toolbox’, with the default alternating least squares algorithm, and random initialization matrices was used.

Images at the soma identified two broad classes of cluster organization identified by NND analyses: one with closely repeating peaks and another with few peaks and greater spatial separation. Long segments of repeating spectrin clusters were particularly evident in neural processes even though isolated clusters were also detected (***Fig. S2D-G***). NND analysis in neural processes confirmed a high proportion of clusters separated with a periodicity of 144 nm, as well as a smaller population of longer inter cluster distances (***Fig. S2D***). Since these patterns at the soma were typically contained within a distance of 1200 nm on NND distributions and ∼150 subpixels on 1D cluster intensity profiles of aligned images, feature categorization of spectrin βII by NNMF in neural processes was restricted to a window of -150 to 100 subpixels (-1128 nm to 752 nm).

NNMF was employed to derive 6 features from the collection of 118 to >2000 (668 ± 25) centroid images stored from each image channel per cell ^25^. Extending the number of features beyond 6 resulted in instances where the centering centroid required as a reference point from which intercluster distances were measured, was diminished or absent in features (***Fig. S3A***). Extracting 6 feature sets from the data reduced this deviation from the true images while still maximizing the number of features to process and the quality of the approximations (***Fig. S3, B and C***). Examples of 6 features extracted from spectrin βIIC labeled neural processes external to the soma of a single cell are shown in ***Fig. S3D***. After smoothing with *spaps*, peaks in the 1D feature data were detected using the *findpeaks* function with a minimum distance, prominence, and height threshold of 12, 0.1% of max, and 5% of max, respectively. 1D features were then classified as regularly spaced clusters (rows) (***Fig. S2F***) when 4 or more peaks were found in the detection window, or isolated clusters when 2 or 3 peaks were located (***Fig. S2G***). Features with less than 2 peaks were discarded.

### Comparison of NNMF to autocorrelation

To fully validate the use of NNMF, we tested its ability to quantify the MPS pattern of spectrin βIIC labeling in neuronal processes compared to conventional autocorrelation (***Fig. S4***). STORM-TIRF images of spectrin in MAP2-positive or MAP2-negative processes outside of the soma provided well defined rows of closely spaced clusters for spectrin βII (***Fig. S4, A and B***). Autocorrelation was applied to 28 of each of these classes of spectrin βII labeled clusters outside of the soma to derive a period of 152 ± 2.38 nm (n = 7) for MAP2 negative processes (***Fig. S4C***). Application of NNMF to MAP2 negative processes calculated a period of 148 nm (R^2^ = 0.9) (***Fig. S4D***). While the period derived by autocorrelation and NNMF are statistically different (p = 0.010, t-stat = 2.6393) the mean values differed by less than 3%. In comparing the results for MAP2 positive datasets, autocorrelation returned a period of 156 ± 4.34 nm and NNMF a period of 157 nm (R^2^ = 0.84), results that were not significantly different (p = 0.752, t-stat = 0.3234) (***Fig. S4, C and D***).

### Fitting of row periods and isolated cluster distances

On the basis of these forms of analyses we applied NNMF to extract features to define the periods of regularly spaced clusters by fitting inter-peak distances with a damped oscillatory model (**Eq. 1**) using *lsqnonlin* in MATLAB (***Fig. S5, A and B***). Fitting this model along the entire data range yielded interpretable results but was optimized to the range of 112-526 nm. The lower limit of this was chosen just within a limit set by the peak detector (90 nm). The higher limit was to ensure the models were fit to the highest signal-to-noise region of the data, and match the most common row length (3 clusters) observed in the soma.

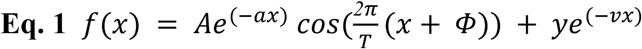

**Equation 1 variables:** A = amplitude; α = amplitude decay; T = period; Φ = horizontal phase shift; y = baseline; v = baseline decay

Optimal fitting was performed on raw and smoothed data (***Fig. S5, A and B***) using preset and user specified initial values for the least squares solver. This raw and smooth data was amplitude normalized prior to initial value selection to make user values more predictable (***Fig. S5B***), but rescaled to the original data after fitting. Four iterative fits were performed using each initial value set with increasing freedom in the parameters (***Fig. S5B***). The parameters that generated the smallest SSQ in the third or fourth fit were used to define the oscillatory model reported.

Isolated features were examined in a window of -376 to -1506 nm (***Fig. S5F***) from the centering cluster to ensure analysis avoided interpretation of very distant clusters and that gaussian fitting was not disrupted by the tails of the centering centroid. Single gaussian fits were applied to data in the examination window using the function *fit*. These gaussian models were used as proxies of the peak data as isolated point-spread-functions are expected to have a gaussian distribution. The standard deviation and mean of these fitted gaussians were plotted on two axes to identify the number of populations in each data set (***Fig. S5G***). Outliers in either standard deviation or mean of the gaussian fit were identified and culled using the function *rmoutliers* in MATLAB (***Fig. S5G***). Remaining gaussian distributions with means outside of the initial examination window (-376 to -1506 nm) or an excessive standard deviation (SD >= 400) were removed. Finally gaussians with a poor goodness of fit (R^2^ ≤ 0.9) were excluded. The remaining mean positions were plotted as a histogram (***Fig. S5H***) and fit with a single gamma distribution with the median reported. The quality of these gamma distributions and their means were assessed by comparing the gamma model median (the median of the means) to the data median along with R^2^.

### Row length and branch angles

To determine row lengths and row branch angles visual counting was performed at high resolution by two viewers. After defining all relevant centroids in the image, a random sample of 5 centroid ROIs per cell were selected with the condition of ROI centers being 600 subpixels apart to prevent double counting. These ROIs were overlaid with detected centroid markers and counted for the length of linear rows with a minimum of 3 clusters drawn from 13-55 cells per label. Up to 388 rows for CaRyK complex proteins, actinin I and II, and spectrin βII were defined across a selection of up to 275 ROIs and 55 cells (**Fig. 6**). A hand selected subset of the randomly picked centroid ROIs was then brightened and used to measure angles when rows of at least 3 consecutive clusters reached a point where a second row of clusters extended with a shift in orientation. Approximately 100 branch points were measured by two viewers for each label across 22-52 ROIs randomly selected from 13-18 cells. Angle measures were generated using Fiji ImageJ.

### Source Data

All 1D NNMF outputs, NND results, fitting results, and representative raw image files have been deposited at the Federated Research Data Repository and are publicly available. Accession numbers and the DOI are located in ***Table S5***. Any further data can be requested from the corresponding author. Original MATLAB and R scripts have been deposited at the Federated Research Data Repository, and are publicly available as of the date of publication.

### Statistical Analysis

NND and NNMF calculations were applied to typically >500 clusters per label/cell (675 ± 26; n = 279) and up to 43,000 clusters in up to 55 cells per label. Smoothing of 1D peak intensity data was done using the spline function *spaps* in MATLAB. Smoothing of raw interpeak data was performed using a moving average *smooth* with its default span of 5 in MATLAB. Interpolation of fitted gamma distributions was utilized to calculate central tendency with higher precision than the bin width of the histogram data. Fitting was performed using two methods. NND distributions and isolated peak positions were fitted with a single or dual gamma distribution using the function *lsqnonlin* in MATLAB. All single distribution fits had their model median compared to the data median to determine general error in this method. Error decreased and R^2^ increased as θ increased. Interpeak distances from row features were fitted with a damped oscillatory model (Eq 1) iteratively using *lsqnonlin* in MATLAB. Goodness of fit was determined with *gfit2* or internally in functions such as *fit.* To determine if distributions were statistically different we utilized the non-parametric Kosmolgov-Smirnov test in MATLAB. To determine if a single value was likely to belong to a distribution a one-sample t-test was used. Comparisons of two normally distributed datasets were determined with two-sample t-tests or Welch’s t-test. Comparisons of centroid counts per image, Mander’s correlation coefficients and overlap coefficients were performed with a Wilcoxon rank-sum test *ranksum* in MATLAB.

## Acknowledgements

We gratefully acknowledge J. Forden for technical assistance, P. Colarusso, C. Brideau, and L. Swift (Live Cell Imaging Facility, Univ. Calgary) for assistance in image calibrations and N.V. Marrion for helpful discussions. This work was supported by the Canadian Institute of Health Research Project Grants MOP136826, PJT 169007 (R.W.T.), Indian Council of Medical Research (ICMR) (IIRP-2023-0253; FIW-2024-01-0000000061; IIRPSG-2025-01-01202) (G.S.), and the Department of Biotechnology (BT/PR47597/BMS/85/46/2024) (G.S.). W.N. holds a Canada Research Chair in Computational Neuroscience

## Author contributions

Authorship: G.S. and D.G. contributed equally to the work; Conceptualization: R.W.T., G.S., W.N.; Methodology: G.S, W.N., D.G., X.Z.; Investigation: G.S., D.G., X.Z., W.N.; Validation: G.S., D.G., W.N., X.Z., R.W.T.; Data analysis: G.S., D.G., W.N., X.Z.; Visualization: G.S., D.G., R.W.T., W.N.; Resources: Live Cell Imaging Facility, University of Calgary; Supervision: R.W.T., W.N.; Funding acquisition: R.W.T., G.S.; Writing, review and editing: All authors wrote or reviewed the manuscript before submission.

## Data and materials availability

All the image data and scripts have been deposited to FRDR (Table S5)

## Supplementary Materials

**Figure S1.**
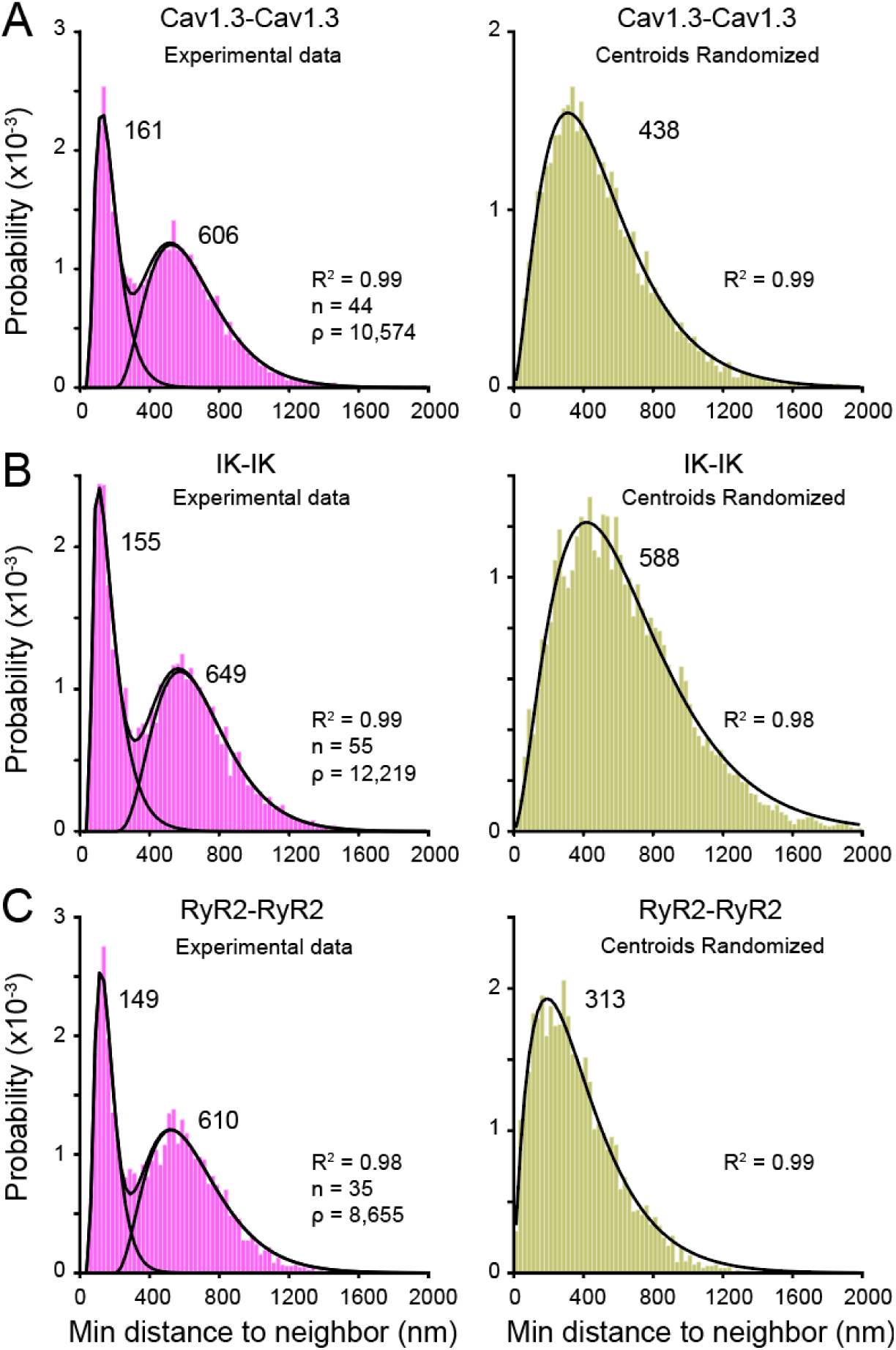
The spatial distribution of CaRyK protein clusters at the soma is dissimilar from a random shuffle. Related to Fig. 1 and 2. (**A-C**) *Left column* provides NND distributions of the indicated CaRyK complex proteins. *Right column* indicates NND distributions for the same datasets after randomizing centroid positions. Significant differences exist between experimental and randomized data for each protein (non-parametric 2- sample Kolmogorov–Smirnov test (KS)): (**A**) *p* < 0.001, KS-stat = 0.1180; (A) *p* < 0.001, KS-stat = 0.1999; (**C**) *p* < 0.001, KS-stat = 0.1957). Numerical indicators adjacent to histograms represent median values. Sample values (n) = cells, ρ, number of centroids detected across all images for a given label set. Bin widths: (**A-C**) 25 nm.

**Figure S2.**
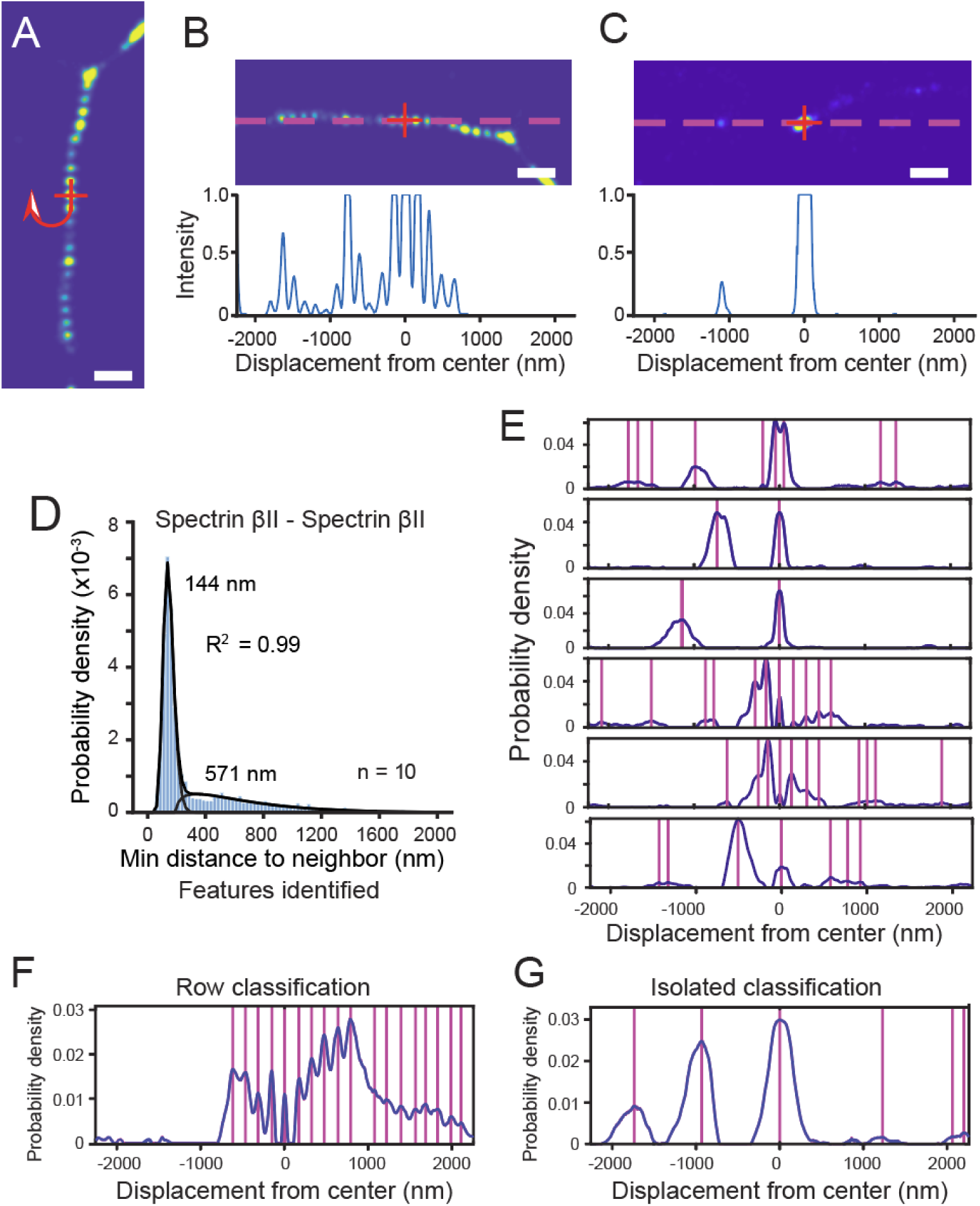
The use of NNMF to define patterns in immunolabeled cluster centroids. (**A-C**) Procedure to define relationships between immunolabeled clusters in representative STORM-TIRF images of neuronal processes outside of the soma labeled for spectrin βII. A systematic pixel-by-pixel search identifies the center of a detected centroid (**A,** *cross*) for spectrin βII labeling. Images are then rotated to a position where its nearest neighboring centroid is oriented to the left and the ROI is stored for feature detection (**B-C**). Below the images in (**B-C**) are 1D displays of normalized pixel intensity aligned with the profile indicated by *dashed lines* for a row of clusters consistent with the MPS (**A-B**) as well as isolated clusters (**C**). (**D**) NND analysis of spectrin βII clusters in neuronal processes reveal two populations of inter cluster distances, with the majority fitting a distribution with a median value of 144 nm. (**E**) Representative example of centroid features extracted by NNMF from spectrin βII labeled neural processes outside the soma of a single cell with k = 6. *Vertical lines* denote detected peak positions. (**F-G**) Two representative examples of features containing rows (**F**) or isolated clusters (**G**) for spectrin βII labeling. Scale bars: (**A-C**) 500 nm. Bin width (**D**) 25 nm.

**Figure S3.**
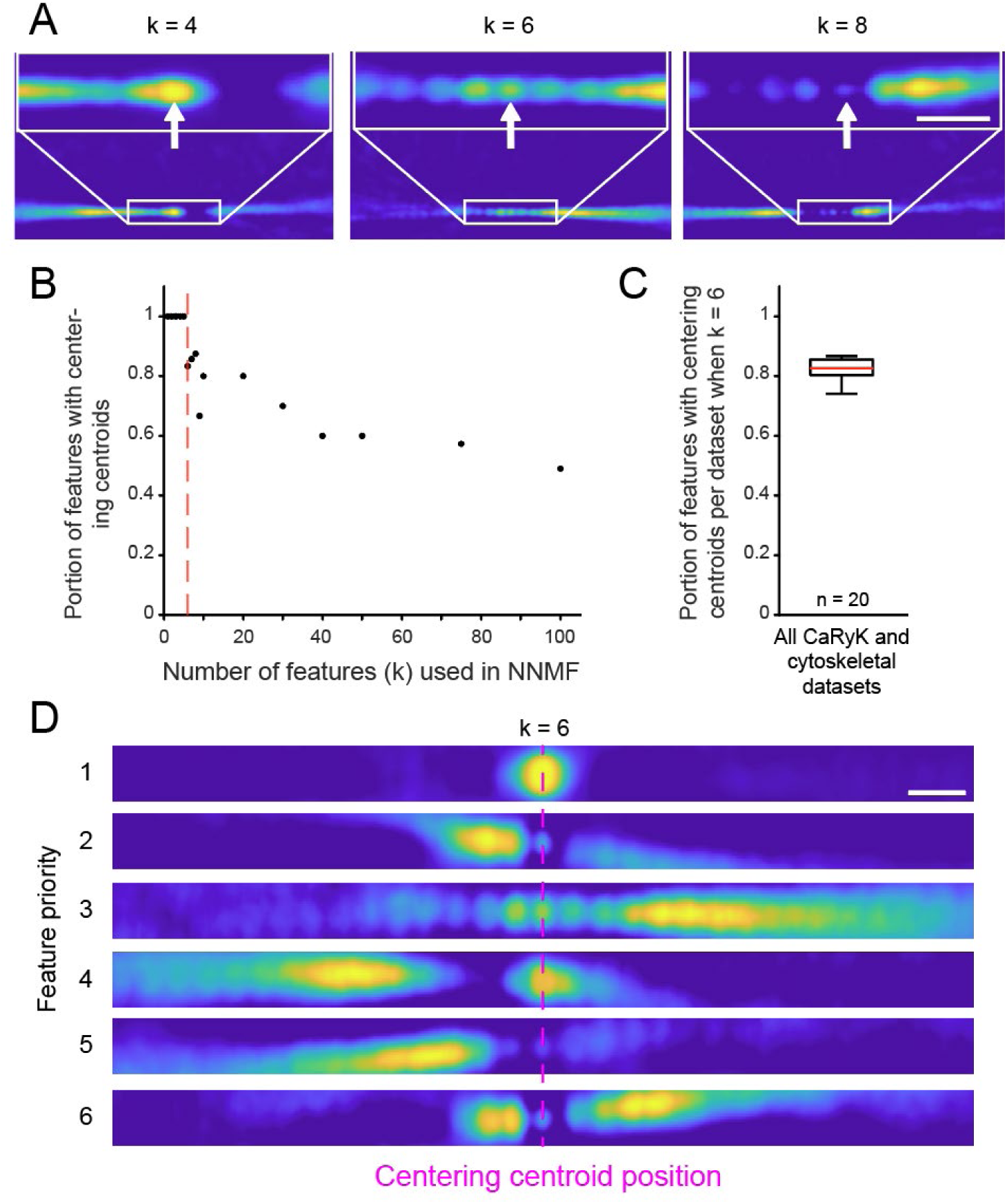
Rationale for feature number selection for NNMF. The effects of using different numbers of features (k) to calculate the distance between rows of clusters using NNMF. NNMF generates features as rows of a matrix which are reported in decreasing importance to the original data set. Every cluster centroid image prior to input into NNMF has a central point spread function upon which the image is aligned to identify intercluster distances of neighboring rows of clusters. (**A**) A representative case of feature detection for different k values for spectrin βII images, with each case illustrating the third feature calculated by NNMF. ROIs of the regions indicated by a *white box* are shown expanded above. As k increases, attributes of each feature become diluted until the central centroid (*white* arrows) becomes diminished or is no more prominent than noise (i.e. k = 8). (**B**) Scatter plot showing a progressive loss of central centroids in downstream analysis as the number of features generated is increased. *Red dashed line* indicates k = 6 (portion = 0.8333). (**C**) Box plot of the portion of features across CaRyK, spectrin and actinin datasets which contain a detectable central centroid to calculate intercluster distances in rows of immunolabeled clusters (mean = 0.824 ± 0.0075). Analyses were conducted with k = 6 to avoid instances where the centering centroid was miniscule or absent and ensure that features reflected a plausible centroid image while maximizing the number of features for downstream analysis. (**D**) Representative example of the top six features extracted for spectrin βII in MAP2 positive or negative processes of hippocampal cells. Scale bars: (**A**, ROIs) 300 nm, (**D**) 300 nm. Sample values (n) in (**C**) reflect all data sets for CaRyK, spectrin and actinin immunolabels.

**Figure S4.**
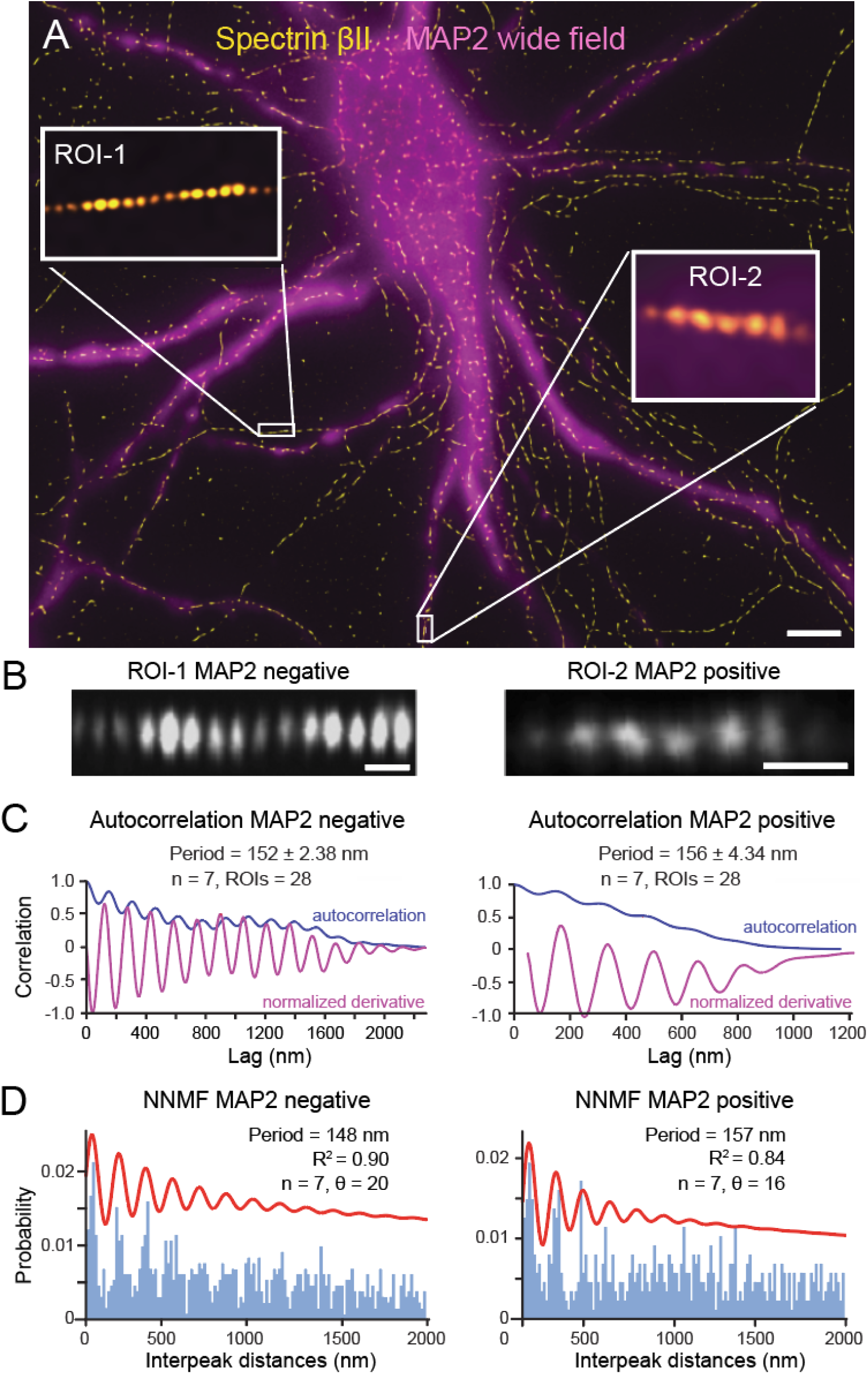
Comparison of periodicity in cluster organization measured by autocorrelation vs NNMF. (**A-B**) Representative STORM-TIRF image of spectrin βII labeling with an overlaid larger field image of MAP2-GFP labeling (**A**). Two processes are highlighted as MAP2 negative putative axons (ROI-1) and MAP2 positive putative dendrites (ROI-2) (**B**). (**C-D**) Spectrin βII labeling of MAP2 negative or MAP2 positive processes outside of the soma are analyzed for periodicity by autocorrelation (**C**) or NNMF (**D**). In (**C**) the autocorrelations (*blue lines*) and normalized derivatives (*magenta lines*) are shown superimposed and drawn from the specific ROI examples in (**B**), while mean periods are calculated from the entire population of clusters. In (**D**) histograms of interpeak distance of row classified spectrin βII features from the same data set are shown superimposed with the damped oscillatory model fits (*red lines*) calculated by NNMF. Scale bars: (**A**) 5 µm, (**B,** ROI-1, 2) 300 nm. Average values are mean ± SEM. Sample values (n) = cells, θ = NNMF row features (**E**). Bin widths: (**D**) 15.4 nm.

**Figure S5.**
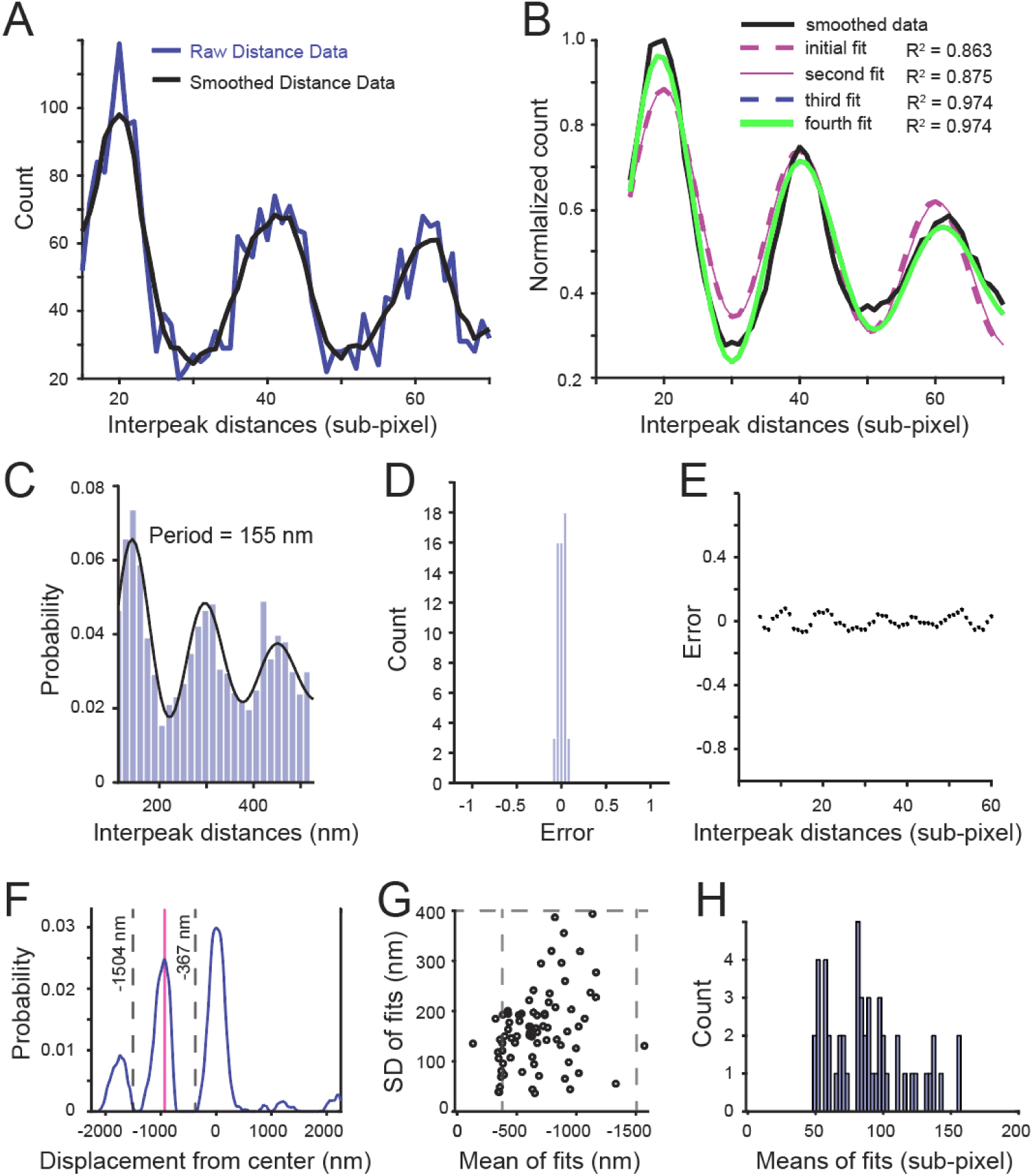
Defining inter cluster distances from NNMF. Procedure for oscillatory model fitting of the NNMF defined interpeak distance histograms. (**A**) Plot of the heights of binned interpeak distances of the initial 3 cycles of row classified features generated by NNMF of spectrin βII labeled processes. Interpeak distances are cropped to a range of 15-70 sub-pixels to accommodate the highest S/N region. Shown superimposed is data before and after smoothing with a moving average to improve model fit. (**B**) Superimposed records of smoothed data trace from (**A**) fit with a damped oscillatory model for 4 iterations using a least-squares method. (**C-E**) Representative example of best fit model and error values for spectrin βII NNMF calculations in processes, with the best fit model (*black line*) superimposed on the histogram of raw interpeak distance data in (**C**). Shown in (**D-E**) is the histogram of error between model and data (**D**), with error distributed around zero. A scatter plot of error along the X axis of data in (**C**) is relatively homoscedastic (**E**). (**F-H**) Procedure for defining peak centers of isolated features identified by NNMF. A representative example of an isolated classified feature as a plot of probability in relation to distance from centering centroid for spectrin βII - spectrin βII (**F**). An examination window of -1504 to -367 nm (*gray dashed lines*) is used to avoid the influence of the centering centroid or distant data tails in defining the peak of isolated clusters (*magenta vertical line*) (**F**). (**G**) Data within the examination window is fit with a single gaussian. A plot of the mean and SD of fits of isolated cluster gaussians is used to confirm a single population of gaussians and eliminate outliers defined using *rmoutliers*. Any gaussians that fell outside of the examination window (*gray dashed lines*) were removed, along with gaussians that were not well fit to the data they represented (R^2^ ≤ 0.9). (**H**) Final histogram of accepted means for isolated cluster distances determined in (**F-G**). Bin widths: (**C**) 15 nm, (**D**) 0.04, (**H**) 20 nm.

**Figure S6.**
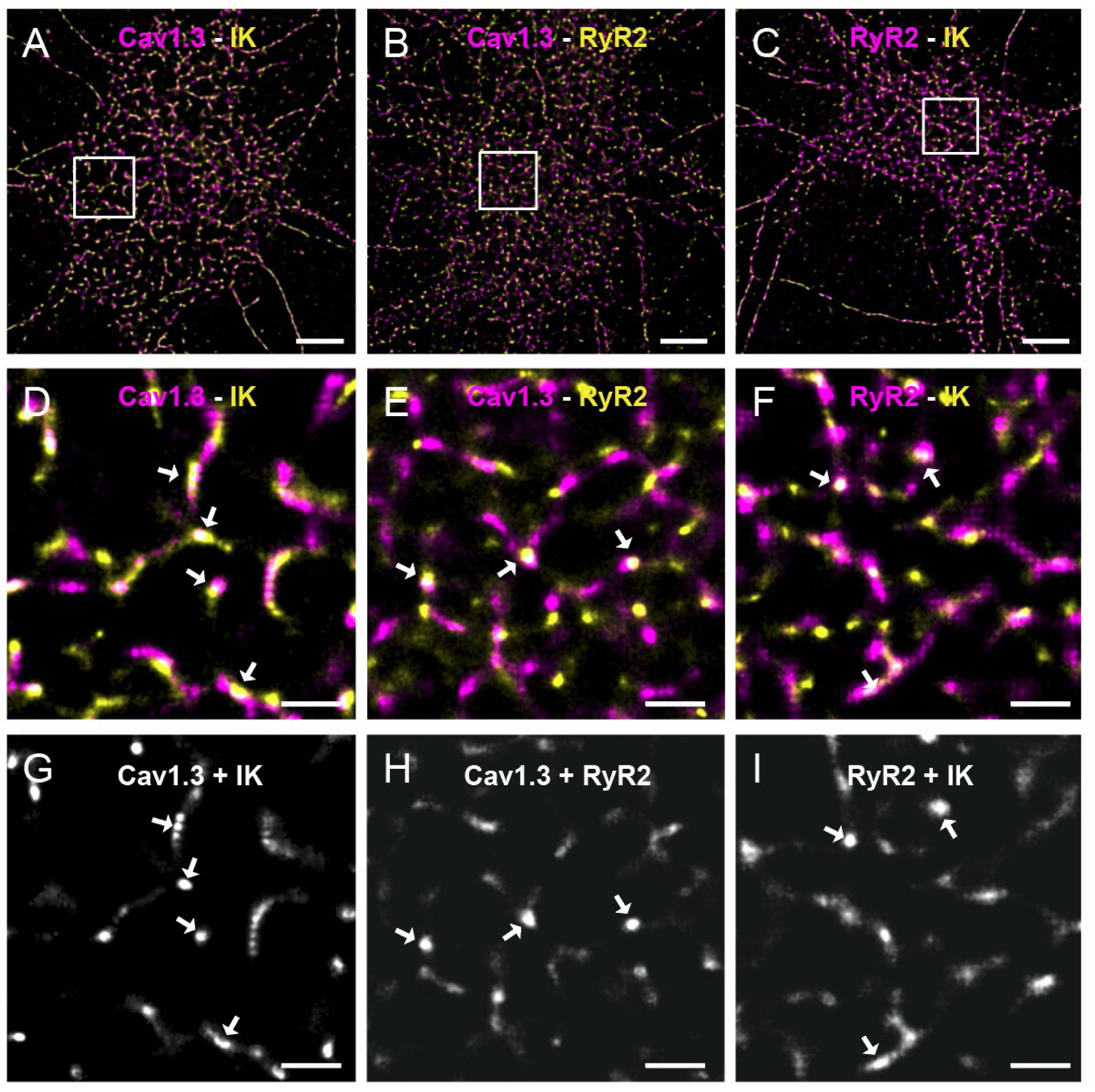
Generating images of fluorescent emission overlap for NNMF analysis. Related to Fig. 1-5. (**A-C**) Larger field views of hippocampal neurons dual labeled for the indicated proteins. (**D-F**) Expanded views of ROIs from images in (**A-C**) illustrating overlap of some fluorescence emissions (white chroma, *arrows*) suggestive of protein colocalizations. (**G-I**) Black and white images of overlap-dependent intensity representations of the dual labels in (**D-F)** to apply NNMF on dual labeled clusters. Pixel intensity of the overlap image is determined pixel wise by 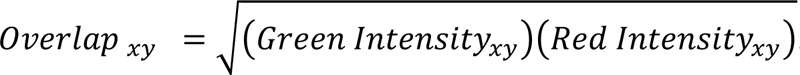. Scale bars: (**A-C**) 5 µm; (**D-I**), 500 nm. See also ***Table S4*.**

**Figure S7.**
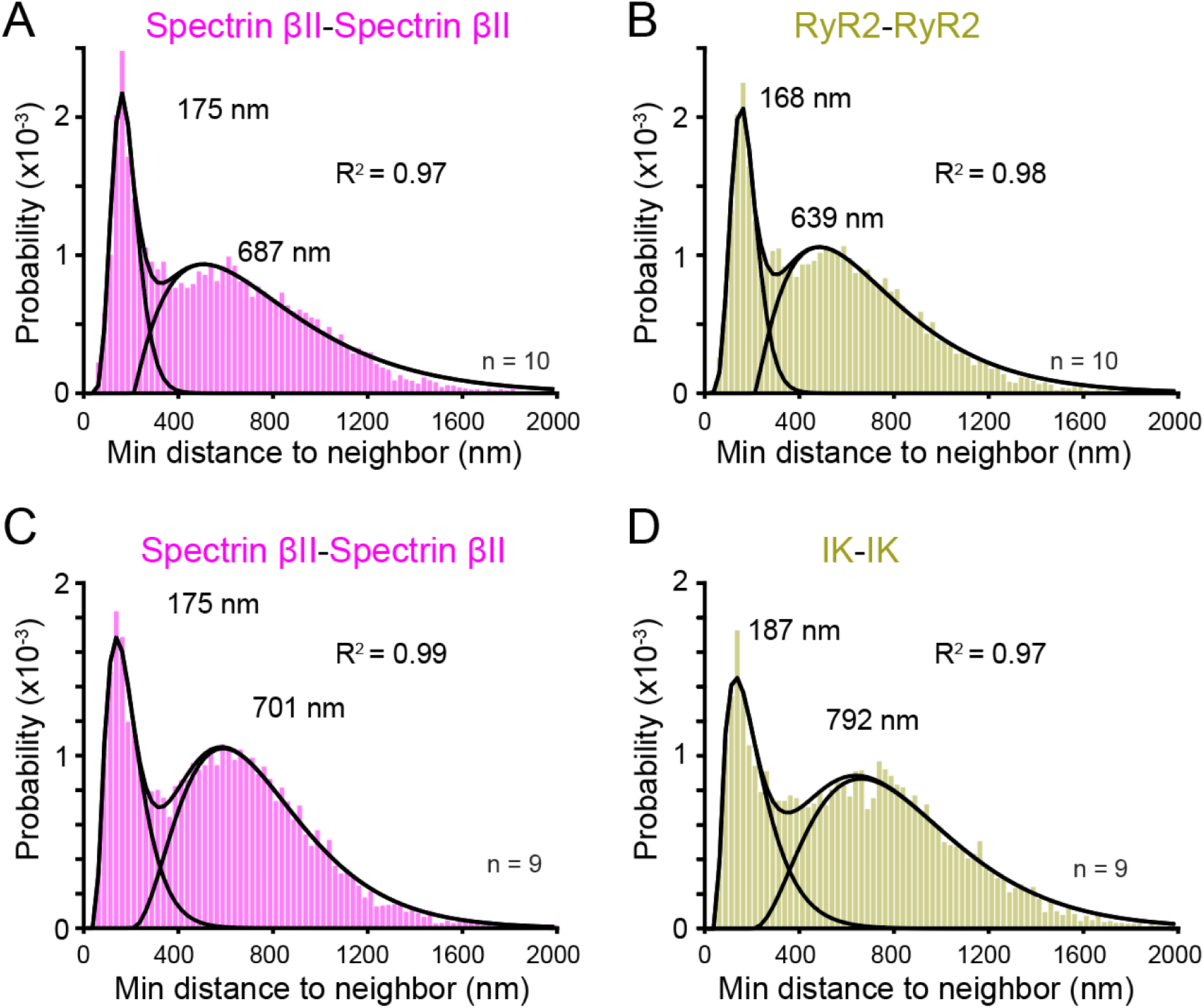
Spectrin βII clusters distribute with similar patterns as IK and RyR2. Related to Fig. 4. (**A-D**) NND histograms of similar immunolabels in two sets of dual labeled tissue (**A-B,** spectrin βII-RyR2; **C-D,** spectrin βII-IK) reveals populations that can be described by two gamma fits. Values are the median of fit gamma distributions and R^2^ of the combined fits. Sample values (n) represent the number of individual cells analyzed. Bin widths: 25 nm.

**Figure S8.**
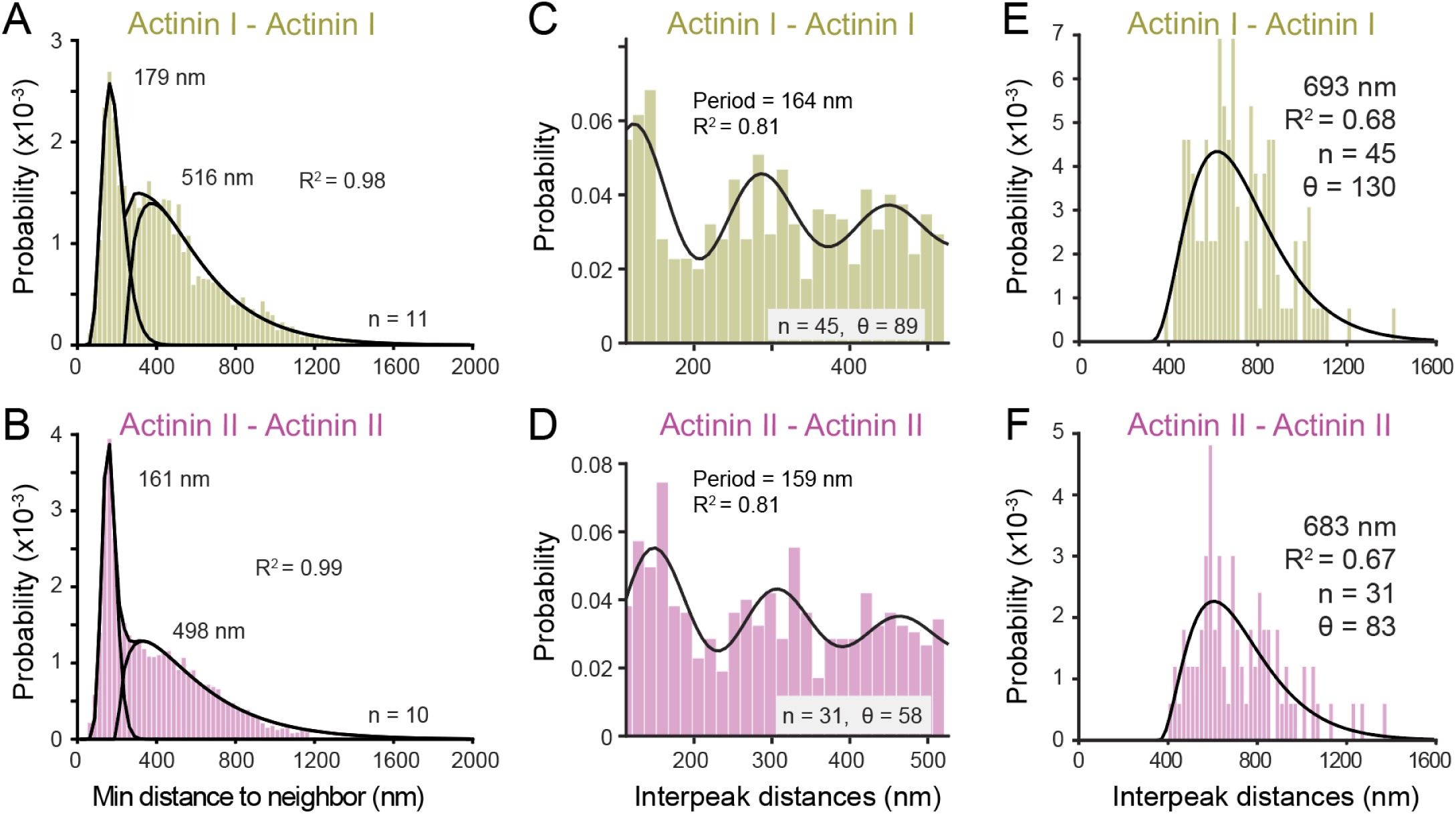
Actinin I and actinin II immunolabels exhibit a structured pattern indicative of rows and isolated clusters. Related to Fig. 5. (**A-B**) NND analysis within actinin I clusters (**A**) and actinin II clusters (**B**) reveal two populations of inter-centroid distance fit with two gamma distributions. Distributions are comparable to that of spectrin βII (see Figs. 3 and 4). (**C-F**) NNMF analysis of the indicated immunolabels defines a regular periodicity of cluster rows (**C-D**) or isolated clusters (**E-F**). Medians of gamma distributions, R^2^ values of fitted gamma distributions (**A, B, E,** and **F**), model periods and with their associated R^2^ values (**C-D**) are indicated on plots. Total number of centroids detected to apply NNMF (**C-F**): actinin I-actinin I, 19,767; actinin II-actinin II, 21,634. Sample values (n) = cells, θ = features. Bin widths: (**A-B)** 25 nm, (**C-D**) 15 nm, (**E-F**) 20 nm. *S1 and S2*.

**Figure S9.**
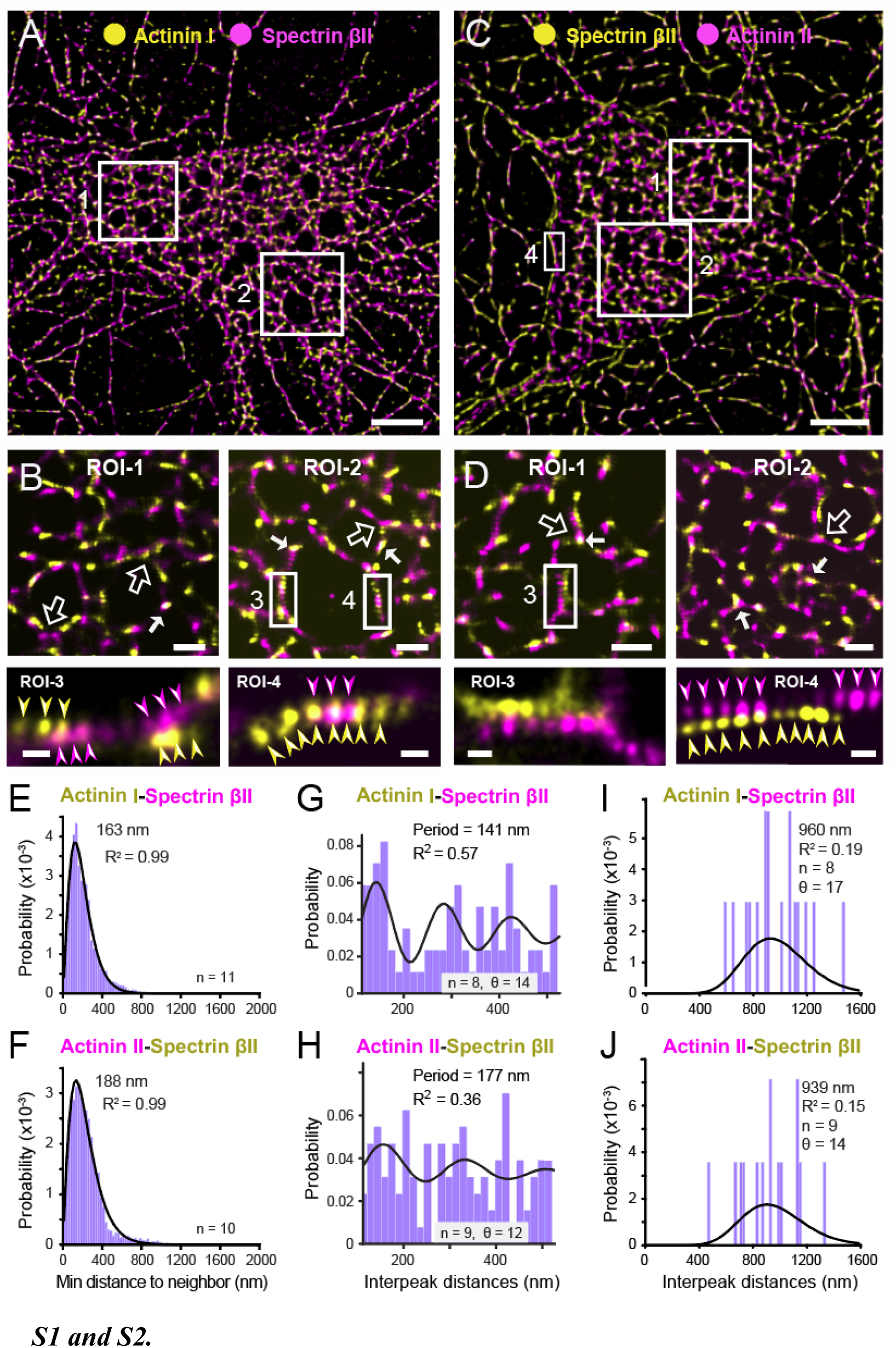
Actinin I and actinin II exhibit an organized pattern in relation to spectrin βII. Related to Figure 5. (**A-D**) STORM-TIRF images of immunolabeled clusters for the indicated proteins. Shown are dual labeled images for spectrin with actinin I (**A-B**) or with actinin II (**C-D**), with ROIs magnified in (**B** and **D**). Rows of spectrin βII or actinin clusters can extend to branch points (*open arrows*), colocalize (*solid arrows*), or alternate to extend or bridge rows of labels (*arrowheads*). (**E-F**) NND analysis between actinin I-spectrin βII (**E**) and actinin II-spectrin βII (**F**). (**G-J**) NNMF analysis defines a periodic pattern for cluster rows (**G-H**) or a single distribution for isolated clusters (**I-J**). Sample values (n) = cells, θ = features. Scale bars: (**A** and **C**) 5 µm; (**B** and **D,** ROI-1, 2), 1 µm; (**B** and **D,** lower ROIs), 200 nm. Bin widths: (**E-F**) 25 nm, (**G-H**) 15 nm, (**I-J**) 20 nm. See also ***Tables***

**Supplemental Table S1.** Summary comparisons of fitted gamma mixture models on NND for the indicated protein pairs. All cells were imaged with two immunolabel targets and NND distributions calculated for all clusters. NND was calculated using LAMA’s morphological cluster analysis and fitted with a mixed gamma model to define central tendency for each gamma distribution in NND histograms. Cluster pairs = the labeled targets which are being compared to generate the NND histogram. n = the number of cells. R^2^ = coefficient of determination when comparing the sum of fitted gamma distributions to the NND histogram. GMM = gamma-mixture model.

| Dual Labeled Tissue | Cluster Pairs | n | First Median (nm) | Second Median (nm) | R<sup>2</sup> | Mixing % of first gamma in GMM |
| --- | --- | --- | --- | --- | --- | --- |
| Cav1.3 - IK | Cav1.3 -Cav1.3 | 11 | 169 | 664 | 0.9773 | 27.2 |
|  | IK-IK |  | 177 | 699 | 0.9827 | 27.7 |
|  | Cav1.3-IK |  | 188 | - | 0.9935 | 100 |
| IK - RyR2 | IK - IK | 11 | 186 | 722 | 0.9822 | 28.1 |
|  | RyR2 - RyR2 |  | 169 | 635 | 0.9888 | 33.2 |
|  | IK - RyR2 |  | 238 | - | 0.9973 | 100 |
| Cav1.3 - RyR2 | Cav1.3 - Cav1.3 | 9 | 174 | 624 | 0.9904 | 34.8 |
|  | RyR2 - RyR2 |  | 185 | 755 | 0.9822 | 25.1 |
|  | Cav1.3 - RyR2 |  | 238 | - | 0.9978 | 100 |
| Cav1.3 - Spectrin βII | Cav1.3 - Cav1.3 | 13 | 168 | 649 | 0.9854 | 30.3 |
| | Spectrin βII - Spectrin βII | | 183 | 733 | 0.9783 | 26.8 |
| | Cav1.3 - Spectrin βII | | 288 | - | 0.9955 | 100 |
| IK - Spectrin βII | IK - IK | 9 | 187 | 792 | 0.9718 | 31.8 |
| | Spectrin βII - Spectrin βII | | 175 | 701 | 0.9879 | 31.1 |
| | IK - Spectrin βII | | 238 | - | 0.9939 | 100 |
| RyR2 - Spectrin βII | RyR2 - RyR2 | 10 | 168 | 639 | 0.9854 | 28.9 |
| | Spectrin βII - Spectrin βII | | 175 | 687 | 0.9720 | 29.4 |
| | RyR2 - Spectrin βII | | 288 | - | 0.9959 | 100 |
| Cav1.3 - Actinin I | Cav1.3 - Cav1.3 | 12 | 180 | 641 | 0.9902 | 23.2 |
|  | Actinin I- Actinin I |  | 192 | 667 | 0.9870 | 18.2 |
|  | Cav1.3 - Actinin I |  | 138 | - | 0.9880 | 100 |
| IK - Actinin I | IK - IK | 11 | 164 | 729 | 0.9872 | 27.7 |
|  | Actinin I - Actinin I |  | 174 | 683 | 0.9757 | 27.6 |
|  | IK - Actinin I |  | 363 | - | 0.9945 | 100 |
| RyR2 - Actinin I | RyR2 - RyR2 | 10 | 168 | 704 | 0.9773 | 24.9 |
|  | Actinin I - Actinin I |  | 179 | 718 | 0.9790 | 21.2 |
|  | RyR2 - Actinin I |  | 338 | - | 0.9931 | 100 |
| Spectrin βII - Actinin I | SpectrinβII - SpectrinβII | 9 | 170 | 489 | 0.9812 | 27.6 |
|  | Actinin I - Actinin I |  | 179 | 516 | 0.9780 | 30.0 |
| | SpectrinβII - Actinin I | | 163 | - | 0.9859 | 100 |
| IK- Actinin II | IK- IK | 11 | 218 | 726 | 0.9800 | 42.5 |
|  | Actinin II- Actinin II |  | 169 | 611 | 0.9901 | 34.2 |
|  | IK- Actinin II |  | 163 | - | 0.9922 | 100 |
| RyR2- Actinin II | RyR2- RyR2 | 10 | 186 | 730 | 0.9799 | 24.9 |
|  | Actinin II - Actinin II |  | 166 | 628 | 0.9822 | 29.1 |
|  | RyR2- Actinin II |  | 213 | - | 0.9966 | 100 |
| Spectrin βII - Actinin II | SpectrinβII - SpectrinβII | 10 | 161 | 498 | 0.9903 | 35.0 |
|  | Actinin II - Actinin II |  | 161 | 452 | 0.9807 | 34.9 |
| | SpectrinβII - Actinin II | | 188 | - | 0.9886 | 100 |

**Supplemental Table S2.** Summary of NNMF feature analysis for the indicated protein pairs. Information about data entered into the NNMF workflow and the results. n = the number of cells imaged. Centroids = the sum of weighted positions across all images that were assessed by NNMF to generate features. Θ = the number of features generated per label set. Row Cluster Data = features from NNMF which were assessed for peak density anticipated in regular and repeating clusters. Isolated Cluster Data = features from NNMF which were assessed with a low peak density anticipated by isolated centroids. Period = period of fitted damped oscillatory model on interpeak data from features. R^2^ = coefficient of determination when comparing data to the fitted model. SSQ = sum of squared error between smoothed interpeak data and the fitted model. Median center = displacement of fitted gamma on isolated features from their reference position. Empty cells represent instances where no interpretable data was produced.

|  |  |  |  | Row Cluster Data |  |  | Isolated Cluster Data |  |
| --- | --- | --- | --- | --- | --- | --- | --- | --- |
| Dataset | n | Centroids<br>$\times 10^3$ | $\theta$ | Period<br>(nm) | $R^2$ | SSQ | Median<br>(nm) | $R^2$ |
| Cav1.3 - Cav1.3 | 45 | 36.8 | 270 | 159 | 0.846 | 0.221 | 638 | 0.733 |
| IK - IK | 53 | 32.7 | 318 | 139 | 0.873 | 0.127 | 714 | 0.653 |
| RyR2 - RyR2 | 55 | 43.1 | 330 | 158 | 0.861 | 0.172 | 711 | 0.543 |
| Spectrin βII - Spectrin βII | 50 | 34.4 | 300 | 159 | 0.920 | 0.121 | 774 | 0.558 |
| Actinin I - Actinin I | 45 | 19.8 | 270 | 164 | 0.810 | 0.230 | 693 | 0.681 |
| Actinin II - Actinin II | 31 | 21.6 | 186 | 159 | 0.813 | 0.225 | 683 | 0.674 |
| Cav1.3 - IK | 11 | 4.1 | 66 | 156 | 0.647 | 0.612 | 795 | 0.308 |
| Cav1.3 - RyR2 | 9 | 3.1 | 54 | 158 | 0.484 | 0.999 | 819 | 0.165 |
| RyR2 - IK | 11 | 4.4 | 66 | 123 | 0.724 | 0.497 | 810 | 0.170 |
| Cav1.3 - Spectrin βII | 13 | 5.6 | 78 | 134 | 0.625 | 0.404 | 917 | 0.165 |
| RyR2 - Spectrin βII | 10 | 3.3 | 60 | 160 | 0.626 | 0.508 | 889 | 0.203 |
| Spectrin βII - IK | 10 | 6.4 | 60 | 121 | 0.509 | 0.710 | 743 | 0.263 |
| Cav1.3 - Actinin I | 12 | 4.2 | 72 | 160 | 0.453 | 1.053 | 775 | 0.396 |
| IK - Actinin I | 11 | 3.5 | 66 | 171 | 0.529 | 0.791 | 906 | 0.141 |
| RyR2 - Actinin I | 14 | 3.1 | 84 | 159 | 0.743 | 0.447 | 868 | 0.382 |
| Spectrin βII - Actinin I | 8 | 6.3 | 48 | 141 | 0.574 | 0.983 | 960 | 0.192 |
| Actinin II - IK | 11 | 7.9 | 66 | 158 | 0.431 | 0.931 | 730 | 0.227 |
| Actinin II - RyR2 | 11 | 4.1 | 66 | 147 | 0.754 | 0.740 | 790 | 0.238 |
| Actinin II - Spectrin βII | 9 | 4.7 | 54 | 177 | 0.356 | 1.027 | 939 | 0.154 |

**Supplemental Table S3.** The distribution of cluster row branchpoint angles are not significantly different. With sample size > 100 normality assumption was ignored and Welch 2 sample t-test was used to compare the central tendency of distributions. Two sample Kolmogorov-smirnov (KS) test was used to test the null hypothesis of the distributions being from the same population. Statistical calculations were completed using R software. df, degrees of freedom.

|  | Welch 2 sample T-test |  |  | KS-test |  |
| --- | --- | --- | --- | --- | --- |
| Angle comparisons | t-stat | df | p-value | k-stat | p-value |
| Cav1.3 - IK | -0.5008 | 197.16 | 0.6171 | 0.0925 | 0.7758 |
| Cav1.3 - RyR2 | 0.2621 | 201 | 0.7935 | 0.0884 | 0.8232 |
| IK - RyR2 | 0.7878 | 194.07 | 0.4318 | 0.1224 | 0.4392 |
| Spectrin βII - Cav1.3 | 0.5033 | 208.84 | 0.6153 | 0.0813 | 0.8836 |
| Spectrin βII - IK | 1.0333 | 196.39 | 0.3027 | 0.1219 | 0.4321 |
| Spectrin βII - RyR2 | 0.2486 | 201.87 | 0.8039 | 0.0577 | 0.9958 |
| Spectrin βII - Actinin I | -1.378 | 214.27 | 0.1697 | 0.1141 | 0.4628 |
| Spectrin βII - Actinin II | -0.2847 | 105.57 | 0.7764 | 0.0962 | 0.8982 |
| Actinin I - Cav1.3 | -0.8699 | 215.32 | 0.3853 | 0.0924 | 0.7313 |
| Actinin I - IK | -0.4210 | 219 | 0.6742 | 0.0947 | 0.7088 |
| Actinin I - RyR2 | -1.149 | 212.22 | 0.2519 | 0.1133 | 0.4852 |
| Actinin II - Cav1.3 | 0.1312 | 102.41 | 0.8959 | 0.0812 | 0.9571 |
| Actinin II - IK | 0.5372 | 88.523 | 0.5924 | 0.0851 | 0.9648 |
| Actinin II - RyR2 | -0.0825 | 100.95 | 0.9344 | 0.1115 | 0.7387 |

**Supplemental Table S4.** Percent overlap of pixels labeled for CaRyK complex proteins. Dual labeled images were masked to their ROIs, thresholded at > 20% maximal intensity and all pixels were counted. When pixels in the same position in both channels were greater than the threshold, they were counted as overlapped and divided by the sum of pixels greater than threshold in each channel. Calculated in MATLAB with custom scripts.

| **A** | **B** | **n** | **% Overlap on (A)** | **% Overlap on (B)** |
| --- | --- | --- | --- | --- |
| Cav1.3 | IK | 11 | 22.5 ± 1.5 | 20.4 ± 1.9 |
| RyR2 | IK | 11 | 13.0 ± 1.8 | 18.2 ± 1.7 |
| Cav1.3 | RyR2 | 9 | 9.6 ± 1.1 | 16.2 ± 1.6 |

**Supplemental Table S5.** List of reagents and supplies used, software applications, and location of data repository.

| REAGENT | SOURCE | IDENTIFIER |
| --- | --- | --- |
| **Primary Antibodies** |  |  |
| Rabbit polyclonal anti-Cav1.3 (1:250) | Alomone Labs | ACC-005; RRID:AB_2039775 |
| Mouse monoclonal anti-Cav1.3 (1:250) | Novus Biol. | NBP2-12893; RRID:AB_2687841 |
| Mouse monoclonal anti-KCa3.1 (1:250) | Santa Cruz | SC-365265; RRID:AB_10841432 |
| Rabbit polyclonal anti-KCNN4 (1:250) | Abcam | ab215990; RRID:AB_3714743 |
| Mouse monoclonal anti-RyR2 (C3-33) (1:200) | Sigma-Aldrich | R-128; RRID:AB_261301 |
| Rabbit polyclonal anti-RyR2 (1:200) | Sigma-Aldrich | SAB4502707; RRID:AB_10746831 |
| Rabbit polyclonal anti-Spectrin βII (1:250) | Abcam | AB264177; RRID:AB_1270902 |
| Mouse monoclonal anti-Spectrin βII (1:250) | BD Transduction | 612563; RRID:AB_399854 |
| Mouse monoclonal anti-Spectrin αII (1:200) | Abcam | Ab11755; RRID:AB_298540 |
| Goat polyclonal anti-Actinin I (1:250) | Novus Biol. | AF8279; RRID:AB_2922129 |
| Rabbit monoclonal anti-Actinin II (1:250) | Abcam | ab72239; RRID:AB_1270902 |
| Rabbit monoclonal anti-Actinin-II (1:250) | Abcam | ab68167; RRID:AB_11157538 |
| Chicken polyclonal anti-MAP2 (1:500) | Abcam | ab5932; RRID:AB_92147 |
| Monoclonal anti-rat CD11b/c microbeads | Miltenyi Biotec | 130-105-634; RRID:AB_2783886 |
| GLAST (ACSA-1)-biotin microbeads | Miltenyi Biotec | 130-095-826; RRID:AB_2733472 |
| **Secondary Antibodies** |  |  |
| Goat F(ab')<sub>2</sub> anti-rabbit IgG–Alexa Flour-647 (1:1000) | Jackson | 111-606-047; RRID:AB_2338082 |
| Rabbit F(ab')<sub>2</sub> anti-mouse IgG–Cy3 (1:1000) | Jackson | 315-166-047; RRID:AB_2340148 |
| Rabbit F(ab')<sub>2</sub> anti-chicken IgG–Alexa Fluor-488 (1:1000) | Jackson | 303-546-003; RRID:AB_2339330 |
| Donkey F(ab') anti-rabbit IgG–Alexa Fluor-64 (1:1000) | Jackson | 711-606-152; RRID:AB_2340625 |
| Goat F(ab') anti-mouse IgG–Cy3 (1:1000) | Jackson | 115-166-146; RRID:AB_2338708 |
| Donkey F(ab') anti-goat IgG–Cy3 (1:1000) | Jackson | 705-166-147; RRID:AB_2340413 |
| Goat anti-chicken IgY (H+L) - Alexa Fluor 488 (1:600) | Abcam | ab150169; RRID:AB_2636803 |
| HRP conjugated goat anti-mouse IgG (H+L) (1:3000) | Invitrogen | 62-6520; RRID:AB_2533947 |
| HRP conjugated donkey anti-rabbit IgG (1:5000) | Cytiva | NA9340; RRID:AB_772191 |
| **Chemicals, peptides, and recombinant proteins** |  |  |
| Isoflurane | Fresenius Kabi | Cat No. CP0406V2 |
| Papain | Worthington | Cat No. LK003150 |
| Dubecco's PBS | Sigma Aldrich | Cat No. D8537-500ML |
| Bovine Serum Albumin | Vector Labs. | Cat No. SP-5050 |
| Normal Goat Serum | Jackson | Cat No. 005-000-121 |
| Glucose oxidase (Aspergellius Niger) | Sigma Aldrich | Cat No. G2133-50KU |
| Catalase (Bovine) | Sigma Aldrich | Cat No. C3155-50MG |
| Beta-Mercaptoethanol | Sigma Aldrich | Cat No. M6250-100ML |
| Triton-X | Sigma Aldrich | Cat No. T-8787 |
| Poly-L-Lysine | Sigma Aldrich | Cat No. P-7280 |
| Neurobasal Medium | Invitrogen | Cat No. 21010-046 |
| Pyruvate | Invitrogen | Cat No. 11360-070 |
| Glutamine | Invitrogen | Cat No. 25030-081 |
| B-27 | Invitrogen | Cat No. 17504-044 |
| Pen-Streptomycin | Invitrogen | Cat No. 15140-122 |
| Laminin | Sigma Aldrich | Cat No. L2020 |
| FBS | Gibco | Cat No. 26140-079 |
| HEPES | Sigma Aldrich | Cat No. H3375 |
| Glucose | Sigma Aldrich | Cat No. G8270 |
| TetraSpek™ beads (100 nm diameter) | ThermoFisher | Cat No. T7279 |
| MACS BSA solution | Miltenyi Biotec | Cat No. 130-091-376 |
| **Other** |  |  |
| No. 1.5 18 mm Coverslip | Electron Microscopy Sci. | Cat No. 72222-01 |
| 70 µm separation filter | Corning | Cat No. 07201431 |
| Minimacs Separator | Miltenyi Biotec | Cat No. 130-042-102 |
| MACS Multi Stand | Miltenyi Biotec | Cat No. 130-042-303 |
| MS Column | Miltenyi Biotec | Cat No. 130-042-201 |
| **Software and algorithms** | Miltenyi Biotec | Cat No. 130-042-201 |
| Fiji | (Schindelin et al., 2012) | https://imagej.net/Fiji |
| LAMA (17.07) | (Malkusch and Heilemann, 2016) | https://www.uni-frankfurt.de/54258347/Software |
| Adobe Illustrator CS6 | Adobe | https://www.adobe.com/ca/products/illustrator.html |
| Matlab (R2022b) | Mathworks | https://www.mathworks.com/products/matlab.html |
| Huygen's Professional (25.04.0p2 64b) | Scientific Volume Imaging | http://svi.nl |
| Origin 8 | Origin Lab | https://www.originlab.com/index.aspx?go=PRODUCTS/Origin |
| **Deposited Data** |  |  |
| MATLAB Scripts | This study | https://doi.org/10.20383/103.01742 |
| Data for MATLAB Scripts | This study | https://doi.org/10.20383/103.01742 |
| **Core Resources** |  |  |
| Live Cell Imaging Laboratory | UCalgary | RRID: SCR_024748 |

